# Detecting chromatin interactions along and between sister chromatids with SisterC

**DOI:** 10.1101/2020.03.10.986208

**Authors:** Marlies E. Oomen, Adam K. Hedger, Jonathan K. Watts, Job Dekker

## Abstract

Accurate chromosome segregation requires chromosome compaction with concordant disentanglement of the two sister chromatids. This process has been studied extensively by microscopy but has remained a challenge for genomic methods, such as Hi-C, because sister chromatids have identical DNA sequences. Here we describe SisterC, a chromosome conformation capture assay that can distinguish interactions between and within sister chromatids. The assay is based on BrdU incorporation during S-phase, which labels the newly replicated strands of the sister chromatids. This is followed by Hi-C, e.g. during different stages of mitosis, and the selective destruction of BrdU containing strands by UV/Hoechst treatment. After PCR amplification and sequencing of the remaining intact strands, this allows for the assignment of Hi-C products as inter- and intra-sister interactions by read orientation. We performed SisterC on mitotically arrested *S. cerevisiae* cells. As expected, we find prominent interactions and alignment of sister chromatids at their centromeres. Along the arms, sister chromatids are less precisely aligned with inter-sister connections every ~35kb. In many instances, inter-sister interactions do not involve the interaction of two identical loci but occur between cohesin binding sites that can be offset by 5 to 25kb. Along sister chromatids, extruding cohesin forms loops up to 50kb. Combined, SisterC allows the observation of the complex interplay between sister chromatid compaction and sister chromatid segregation as the cell transitions from late S-phase to mitosis. SisterC should be applicable to study mitotic events in a wide range of organisms and cell types.

## Introduction

During S-phase, when sister chromatids are formed, they are closely cohesed by the cohesin complex. Sister chromatids are initially also thought to be wrapped around each other and entangled. During the subsequent mitosis, sister chromatids become compacted and, in the process, become disentangled and segregated from each other, although they remain aligned side by side^1^. Classically, this process has been studied using microscopic methods by labeling sister chromatids differently using thymidine analogues^2,3^. It has been difficult to study this complex series of mitotic events using genomic techniques such as Hi-C, as sequencing-based methods cannot distinguish the identical sister chromatids, and therefore cannot differentiate interactions between and along sister chromatids. Recently an assay detecting sister chromatid exchange events allowed mapping of sister chromatid interactions genome wide in bacteria^4^. However, this approach requires extensive genome editing to introduce sister chromatid exchange markers throughout the genome.

Here we present a Hi-C-based assay, SisterC, that can detect and distinguish inter- and intra-sister interactions. We demonstrate the performance of the assay by studying mitotic *S. cerevisiae* cells. In *S. cerevisiae* the cohesin complex mediates inter-sister interactions at the centromere and along the chromosome arms (“cohesive cohesin”)^5,6^. In addition, the cohesin complex compacts the sister chromatid arms by forming intra-sister loops by dynamic loop extrusion (“extruding cohesin”)^7–11^. The latter is different from many other organisms, including mammals, where the condensin complex compacts chromosome arms. Condensin in yeast does act on the centromeres and rDNA loci^7^. SisterC reveals the alignment of sister chromatids, the positioning of inter-sister and intra-sister interactions and how the interplay between cohesive and extruding cohesin shapes the mitotic chromosome.

## Results

### SisterC library production after induction of single strand breaks at BrdU incorporation sites

Sister chromatids are identical in sequence but differ in which strand was newly replicated in S-phase. This difference can be leveraged to differentiate interactions between and within sister chromatids. BrdU containing DNA strands can be selectively degraded after Hoechst treatment and radiation with UV^3,12^. This property has been used before in strand-seq, which allows the detection of sister chromatid exchange events^13–15^. Here we describe SisterC, a method that combines Hi-C^16,17^ on BrdU incorporated DNA and UV/Hoechst treatment to distinguish interactions between sister chromatids (inter-sister interactions) and along sister chromatids (intra-sister interactions). Briefly, SisterC works as follows (figure 1a-c). When cells go through S-phase in the presence of BrdU, this results in pairs of sister DNA molecules containing BrdU in the – strand (assigned as sister A), and DNA molecules with BrdU in the + strand (sister B) (figure 1a). Hi-C ligation of crosslinked and digested fragments of these DNA molecules results in a matrix of 16 possible ligation products between and within sister A and sister B that differ in the orientation of ligated fragments and the strand or strands that contain BrdU (figure 1b-c). Treatment of the ligation products with Hoechst and UV light creates single strand nicks in strands containing BrdU. Upon PCR amplification, this results in specific depletion of Hi-C products for which BrdU was present in both strands and only 8 possible ligation products will be amplifiable (figure 1c). Four of these amplifiable ligation products will be interactions between sister chromatids and four products will be interactions along a sister chromatid, but they differ in fragment orientation. Fragment orientation can be inferred after paired-end sequencing of the SisterC library: + + or − − read orientations represent inter-sister interactions, while + − or − + read orientations represent intra-sister interactions.

**Figure 1.**
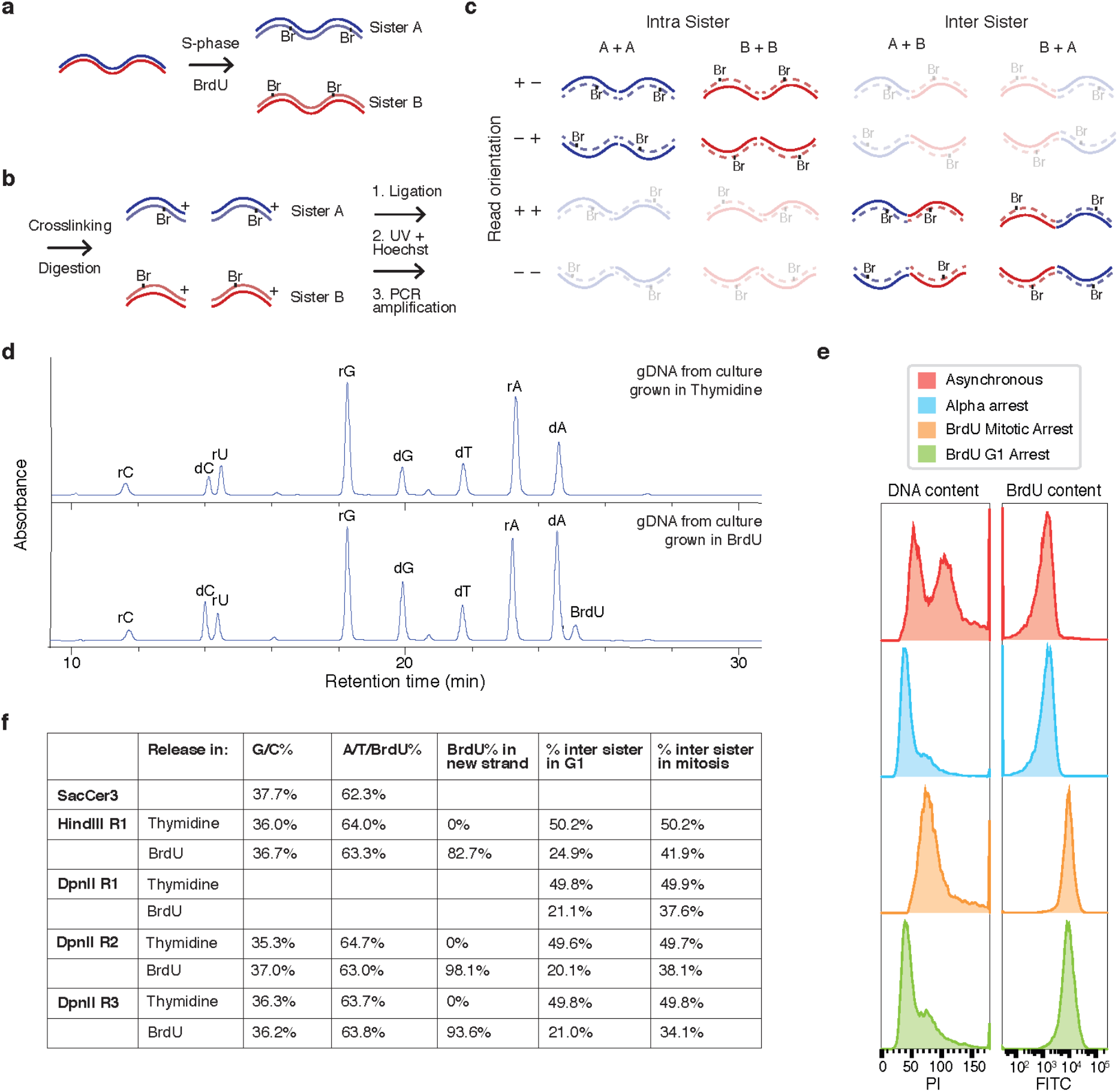
Outline of SisterC chromosome conformation caption technique. **(a)** As cells go through replication BrdU is incorporated on the − strand for sister A and on the + strand for sister B. (**b**) During Hi-C/SisterC preparation DNA is digested followed by proximity ligation, UV-Hoechst treatment and PCR-amplification. (**c**) Proximity ligation leads to 16 possible Hi-C fragments of inter- and intra-sister interactions. For SisterC libraries, DNA molecules were treated with Hoechst and radiated with UV, which introduces single strand nicks. This causes a depletion of DNA strands containing BrdU. Therefore, we can identify inter-sister (+ + and − − reads) and intra-sister interactions (− + and + −) by read orientation. (**d**) HPLC spectra of DNA from yeast cultures released in BrdU and Thymidine allow detection and quantification of BrdU content. Percentages of each nucleoside are then calculated using the extinction coefficient of each nucleoside (**e**) Flow cytometry analysis of cell cycle profile and BrdU incorporation of harvested yeast cultures for preparation of SisterC libraries. (**f**) Table of BrdU content and percentage inter-sister interactions in each SisterC replicate preformed in this study.

Here we choose to study mitotic sister chromatid interactions in *S. cerevisiae* using SisterC. Budding yeast has a relatively small and haploid genome. Furthermore, it can be synchronized in both late G1 and mitosis, which allow to controlled incorporation of BrdU for exactly one S-phase. Lastly, its mappable centromeres and known cohesin binding sites along arms offer sites of particular interest, as these are sites at which sister chromatids are connected and intra-sister chromatid loops may form. As wild-type *S. cerevisiae* cells do not take up nucleosides from the environment, we use a strain that expresses human equilibrative nucleoside transporter (hENT) and *Drosophila* deoxyribonucleoside kinase DmdNK, which allows cells to take up BrdU from the environment and incorporate it into their DNA^18^. Cells were synchronized in late G1 using alpha factor, the culture was split and released in media containing either BrdU or thymidine. This was followed by nocodazole treatment to obtain cells arrested in mitosis or was followed by a second alpha factor incubation to obtain cells arrested in the subsequent G1 (supplemental figure 1).

To investigate the efficiency of SisterC, we performed several control experiments. First, to determine the amount of BrdU incorporation required to achieve efficient Hoechst/UV-induced strand destruction, we PCR amplified DNA fragments using a range of dTTP to BrdUTP ratios, resulting in products with BrdU incorporated in both strands. After treatment with Hoechst and UV radiation, we amplified these products again (supplemental figure 2). This showed that PCR amplification efficiency of DNA fragments containing more than 10 to 50% BrdUTP after treatment with both UV and Hoechst is greatly reduced, indicating that template strands were successfully broken. Second, we performed flow cytometry to detect BrdU incorporation and cell cycle profile of the synchronized yeast cultures (figure 1e and supplemental figure 3). We observe proper cell synchronization in mitosis and late G1 and uniform BrdU incorporation across the cell population. Third, we directly measured the BrdU incorporation efficiency in yeast cells by determining the base composition of genomic DNA using HPLC. HPLC allows to quantitatively measure each nucleotide present in genomic DNA extracted from cultures grown in BrdU or thymidine (figure 1d and supplemental figure 4). We identified all peaks in the HPLC spectra by mass spectrometry (supplemental figure 5) and detected all DNA and RNA nucleosides, including BrdU. After adjustment of the peak area in the HPLC spectra by extinction coefficient, we find that 82.7 up to 98.1% of all Ts in the newly replicated strand are replaced by BrdU (figure 1f and supplemental table 1). Lastly, we estimated the efficiency of selective depletion of Hi-C ligation products as result of UV/Hoechst treatment by producing SisterC libraries for cells synchronized in late G1 after BrdU incorporation during the previous S-phase. As each cell went through the cell cycle and divided into two cells, there are no longer sister chromatids in G1 cells. Using these cells to perform SisterC, we were able to estimate the percentage of false inter-sister interactions as these should no longer be present in a G1 SisterC library. Although we do not detect full depletion of all inter-sister interactions in G1 libraries, we do see a depletion down to approximately 20% (figure 1f and supplemental table 1). In contrast, in mitotic SisterC libraries we typically see around 35-40% captured inter-sister interactions (figure 1d and supplemental table 1), supporting that in SisterC inter-sister interactions are enriched in pairs with − − and + + read orientation and intra-sister interactions are enriched in pairs with − + and + − read orientation.

**Figure 2.**
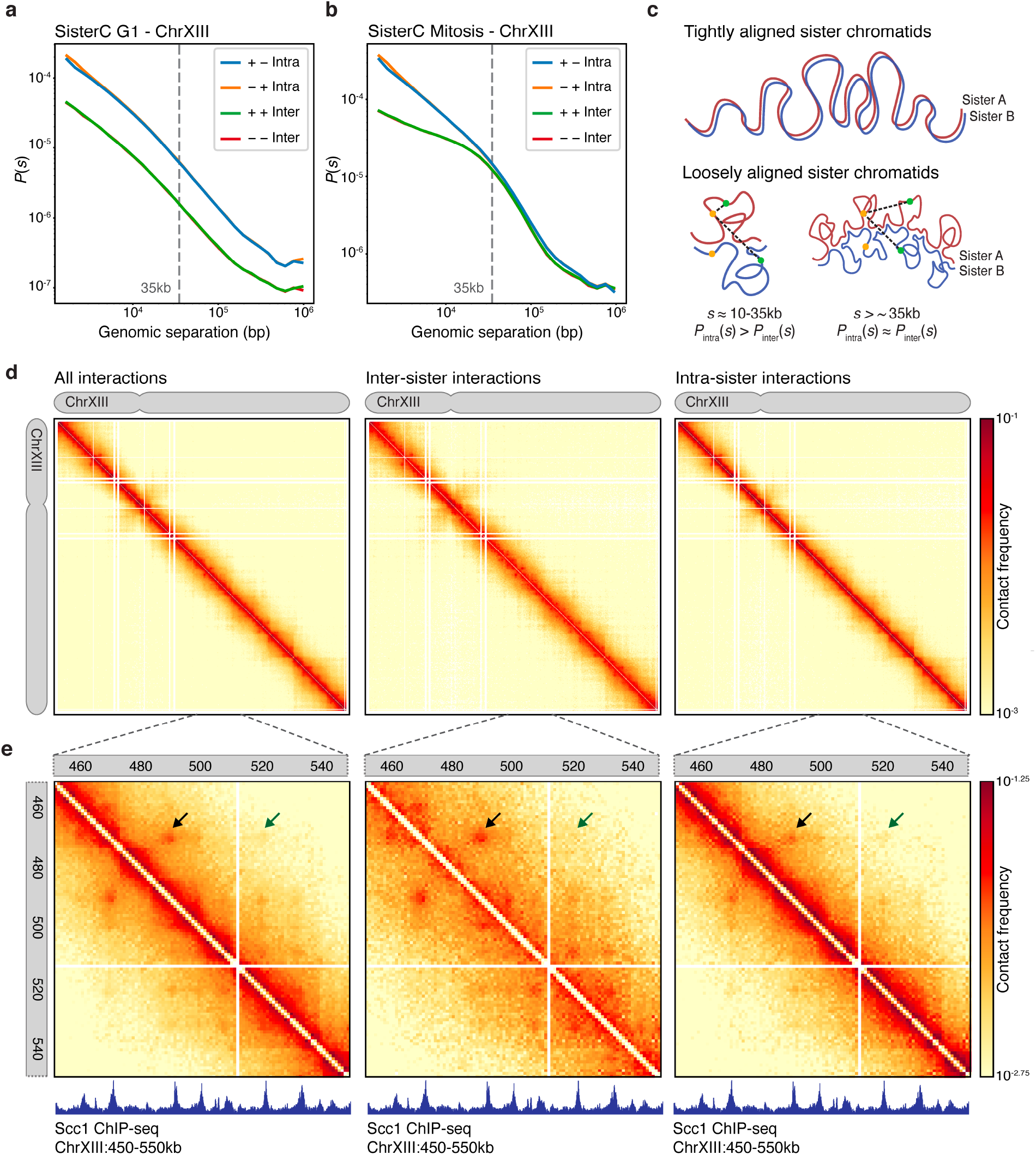
SisterC allows differentiation of inter-sister and intra-sister interactions in mitotic chromosomes. **(a)** Distance decay show uniform depletion of reads with + + and − − orientation in G1 synchronized yeast of fragments with larger than 1500bp genomic separation. (**b**) Mitotic SisterC libraries show an offset of inter-sister interactions up to 35kb genomic separation. (**c**) Model of tightly aligned sister chromatids (top panel) and loosely aligned sister chromatids (bottom panel). (**d**) Chromosome-wide SisterC interactions on chrXIII of all reads (left panel), all reads assigned as inter-sister interactions (middle panel) and all reads assigned as intra-sister interactions (right panel), binned in 2kb bins. (**e**) SisterC interactions at 1kb resolution of all reads on zoomed in region of chrXIII:450,000-550,000 of all reads (left panel), inter-sister reads (middle panel) and intra-sister reads (right panel). Arrows highlight interaction of cohesin sites that are more prevalent as inter-sister interaction. Lower panels show Scc1 ChIP-seq track, a subunit of the cohesin complex.

**Figure 3.**
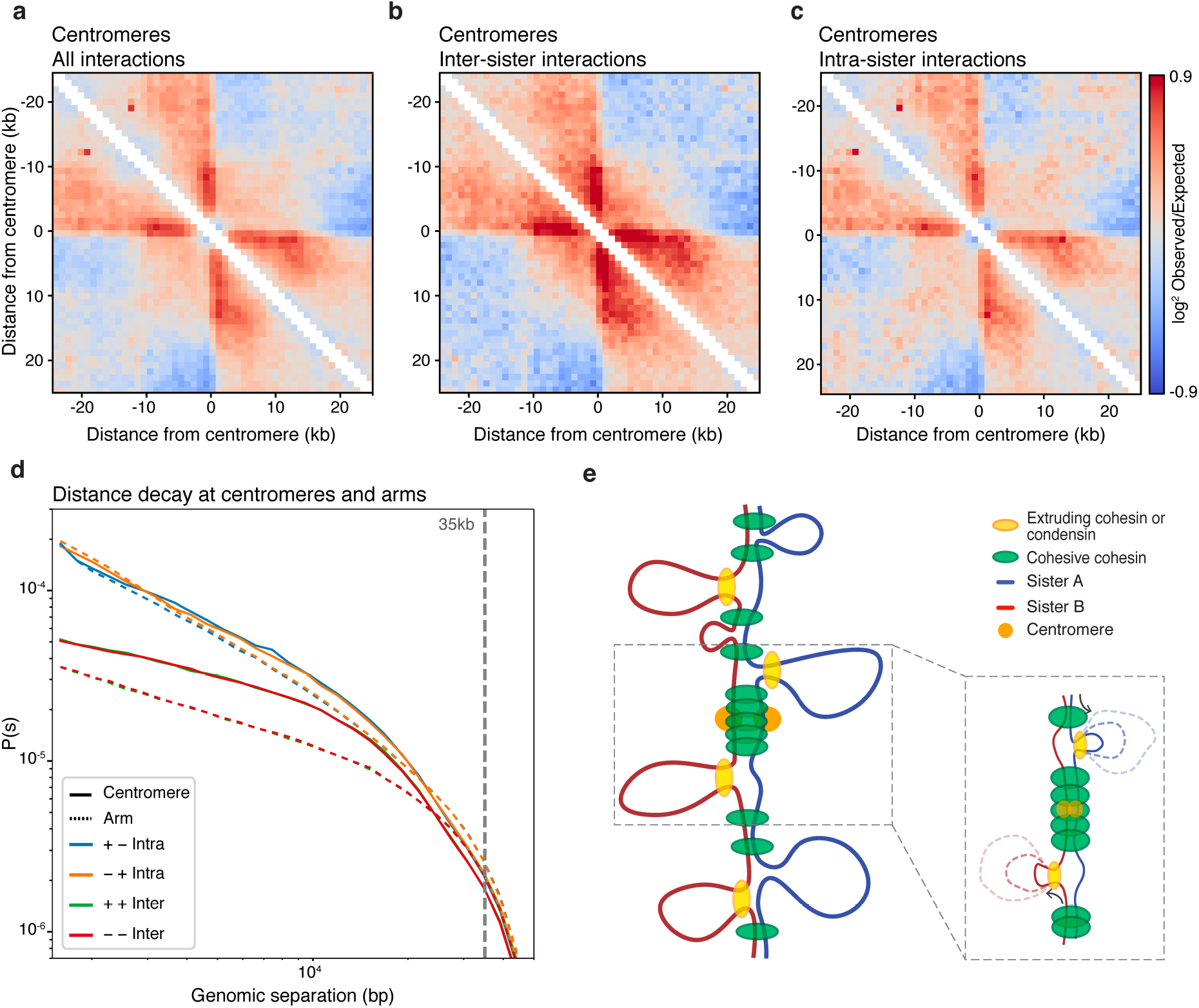
SisterC data show different conformation state at centromeres compared to chromosome arms. (**a-c**) Pile up plot on all yeast centromeres at 1kb resolution of all interactions (**a**), inter-sister interactions (**b**) and intra-sister interactions (**c**). (**d**) Distance decay show a less distinct offset of inter-sister interactions at centromeric regions (solid lines) compared to regions along the chromosome arms (dashed lines). (**e**) Proposed model of centromeric sister chromatid conformation where a condensed array of loops mediated by cohesin and condensing molecules mediate tight and aligned interaction of the sister chromatids at the centromere. Away from the centromere a less dense loop array of cohesin loops result in an offset of inter-sister interactions. As centromeres function as a fixed loop anchor, intra-sister loops can only be formed emerging from one direction (see insert), as observed in a-c.

**Figure 4.**
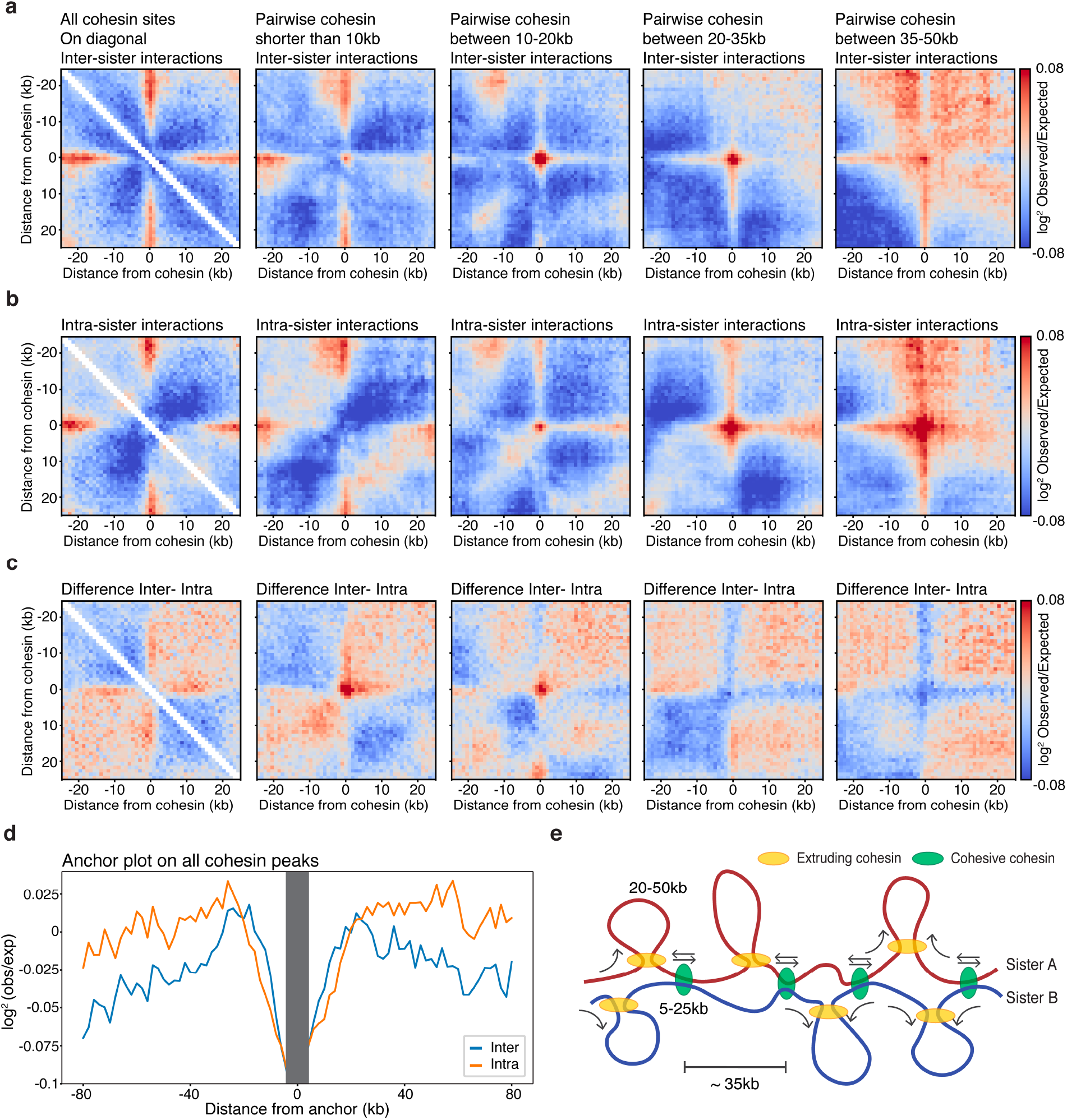
SisterC shows that cohesive cohesin mediates interactions between sister chromatids at shorter genomic distances than loops formed within sister chromatids by extruding cohesin. (**a-b**) Pile up plot of inter sister interactions (**a**), intra-sister interactions (**b**) and the difference between inter-sister and intra-sister interactions (**c**) of all cohesin sites (left panel), of all pairwise cohesin interactions shorter than 10kb (second to left panel), all pairwise interactions of cohesin sites between 10-20kb apart (middle panel), all pairwise interactions of cohesin sites between 20-35kb apart (second to right panel), and cohesin sites between 35-50kb apart (far right panel). (**d**) Anchor plot of all cohesin binding sites show that inter-sister interactions are preferentially formed at distances shorter than 30kb, whereas intra-sister interactions at cohesin sites predominantly interact with sites further than 25kb away. (**e**) Proposed model of intra-sister interactions formed by extruding cohesin and inter-sister interactions formed by cohesive cohesin. Due to a sparse cohesin array, loops of different sizes are made within sisters by extruding cohesin, which results in an offset of 5-25kb in interactions between sister chromatids mediated by cohesive cohesin. The average distance between two cohesive cohesins is 35kb. Intra-sister interactions mediated by extruding cohesin are typically 20 to 50kb in size.

### SisterC allows observation of interactions between and along sister chromatids

We carried out three biological replicates of SisterC experiments using DpnII with highly concordant results and one SisterC experiment using HindIII (supplemental figure 6 and supplemental table 1). After mapping and standard Hi-C data processing (see methods), we examined the SisterC interaction frequency (*P*) as a function of genomic distance (*s*) separated by strand orientation (figure 2a-b and supplemental figure 7a-b). Here the definition of genomic distance for interactions between sister chromatids is the difference between their respective genomic coordinates, even though this involves two different DNA molecules. We compared the results to Hi-C controls (supplemental figure 7c-d) and SisterC negative controls of cells cultured in thymidine instead of BrdU (supplemental figure 7e-h). As expected from any Hi-C library, read orientation of interactions shorter than 1500bp are influenced by technical artifacts, such as unligated dangling ends (+ − orientation) and self-ligated fragments (− + orientation)^19^. Therefore, we exclude any interactions between loci separated by less than 1500bp in all our analyses (supplemental figure 7a-b). In the G1 SisterC libraries (figure 2a), we see a clear depletion in *P*_inter_(*s*) of inter-sister interactions as expected, as there are no longer sister chromatids in G1. Note that the *P*_inter_(*s*) of interactions assigned as inter-sister contacts runs parallel to interactions assigned as intra-sister. This suggests that the former interactions with + + and − − read orientations are in fact intra-sister interactions that were not successfully depleted by UV/Hoechst treatment.

Interestingly, we find a very different *P*_inter_(*s*) for inter-sister interactions in mitosis (figure 2b). Inter-sister interactions are less frequent than intra-sister interactions on short distances (below 30kb), however *P*(*s*) for inter-sister and intra-sister interactions converge for distances larger than 35kb. This describes two phenomena. First, sister chromatids in yeast are not perfectly aligned, as perfectly aligned sister chromatids would result in a minimal difference in *P*(*s*) between inter-sister and intra-sister interactions (figure 2c, top model). Instead interactions with genomic distance below 35kb occur more frequently within the same sister than interactions between sisters, suggesting sister chromatids are loosely aligned (figure 2c, bottom model, left panel). Second, interaction frequency of inter-sister and intra-sister interactions between loci separated by more than 35kb distance converge, which implies that the likelihood of an interaction being inter-sister or intra-sister has become identical (figure 2c, bottom model, right panel). Together, this suggests that although the sister chromatids are not perfectly aligned, sisters are loosely held together mediated by inter-sister interactions that are spaced every 35kb on average (figure 2c).

When mitotic SisterC data are visualized as an interaction heatmap, e.g. for chromosome XIII (figure 2d), we observe similar characteristics as seen in *P*_inter_(*s*). Inter-sister interactions show a weaker interaction signal around the diagonal, representing short-range interactions up to 30kb, compared to intra-sister interactions. At larger distances this difference in interaction frequency is no longer detectable. However, zooming in to a smaller region on chromosome XIII, we observe features that appear stronger in either the inter-sister heatmap or the intra-sister heatmap (figure 2e). In the combined SisterC interaction map (the sum of inter- and intra-sister interactions, figure 2e left panel), we observe dots that represent interactions between cohesin binding sites as detected by ChIP-seq^20^. When inter-sister or intra-sister interactions are plotted separately, we observe that some of these interactions are more prominent in the inter-sister dataset (black arrow, figure 2e), while others are more prominent in the intra-sister dataset (green arrow, figure 2e). This is difference is further highlighted after correction for the distance dependent expected interaction frequency (supplemental figure 8). The finding that some of these interactions between cohesin binding sites are detected in both inter- and intra-sister datasets, albeit with different frequencies, could be related to the fact that Hoechst/UV depletion is not complete. Perhaps more interestingly, this could also be due to variability in the population in the positioning of inter- and intra-sister interactions. Where in one cell a given cohesin site interacts with a second cohesin site along the same chromatid to mediate an intra-sister interaction, in another cell this cohesin site interacts with the same second cohesin site but located on the other sister chromatid to form an inter-sister interaction. Figure 2e illustrates a second example of cell to cell variability: the inter-sister interactions highlighted by the black arrow and the intra-sister interaction highlighted with the green arrow, are mediated by the same cohesin binding site at position chrXIII:469,450. One interpretation is that in one cell this site is involved in an inter-sister interaction where in another cell this site is mediating an intra-sister interaction. Interestingly, inter-sister interactions seem to occur at a shorter genomic distance than intra-sister interactions, as will be described in more detail below.

### Sister chromatid interactions at centromeres

We set out to explore SisterC data around centromeres as we expect enrichment of inter-sister interactions at these sites. Centromeres display very prominent binding of both condensin and cohesin, where they mediate inter-sister interactions and possibly intra-sister interactions^7^. When we aggregate all inter- and intra-sister interactions combined around all 16 centromeres (figure 3a), we observe a striking pattern. Regions directly adjacent on either side of the centromere interact frequently with sequences up to 10 to 15kb away from the centromere. When interactions between sisters and interactions along the sisters are plotted separately (figure 3b-c), we find that inter-sister interactions contribute differently to the combined heatmap than intra-sister interactions do. First, intra-sister interactions are depleted in a 2kb window centered precisely at the centromere. Second, we observe that, compared to the genome wide average expected level, inter-sister interactions at centromeric regions are enriched up to 10kb away from the diagonal (figure 3b and 3d). Third, both intra- and inter-sister interactions contribute to the line-like features emanating from the centromere in the Hi-C maps. Line-like features in Hi-C maps have been interpreted as dynamic or variable loops with one fixed anchor and with the second anchor at various distances away^21^. In this instance the fixed inter-sister connection and intra-sister loop anchor is located directly adjacent to the centromere. This fixed site then is engaged in an inter-sister interaction with a site located on the other sister chromatid some variable distance away from the centromere but on the same chromosome arm. Similarly, within a sister chromatid the fixed loop anchor would engage with a second anchor at some variable distance. Averaged over 16 centromeres, lines appear, but at individual centromeres such intra-sister looping and inter-sister interactions at the same site can be observed as an enriched dot, an example is shown in supplemental figure 9.

As described above, we observe that on average *P*_inter_(*s*) is smaller than *P*_intra_(*s*) for loci separated for up to 35kb, while they converge for loci separated by larger distances. When we plot these two parameters for a 50kb window around all centromeres (figure 3d, solid lines) and compare it to a similar sized window on the chromosome arms (figure 3d, dashed lines), we observe a much smaller depletion of *P*_inter_(*s*) for short range interactions. Additionally, around centromeres *P*_inter_(*s*) and *P*_intra_(*s*) converge around *s* = 15kb. This indicates that around centromeres sister chromatids are more closely aligned at least in part because cohesin mediated sister interactions are spaced more closely: spacing between ChIP-seq cohesin peaks is about 5kb around centromeres and the peaks are much higher than along arms, indicating that every cell obtains an array of closely spaced bound cohesin at centromeric regions during mitosis^20,22^.

Combined these observations lead us to propose that around centromeres interactions between and along sister chromatids are organized differently from the chromosome arms. More frequent binding of cohesin results both a denser loop array and a tighter alignment of the sister chromatids. We note that we cannot differentiate between a situation where extruding loops are actively formed by a loop extruding factors such as extruding cohesin and condensin or if the observation of loops in centromeric regions is a result of offset cohesive cohesin binding, which would passively expulse a loop on one sister chromatid (figure 3e). Both scenarios will be observed as line-like features in Hi-C heatmaps.

### Inter-sister and intra-sister interactions along arms are mediated by independent cohesin complexes that act at different genomic distances

Cohesin mediated interactions along chromosome arms have been identified by Hi-C before^7,23,24^, however prior to SisterC it has not been possible to differentiate between interactions between and along sister chromatids. When we plot all inter-sister interactions (figure 4a) and intra-sister interactions (figure 4b) at and around individual cohesin sites (far left panels), we see that cohesin sites preferentially interact with sites located at least 5kb away on either side. Interestingly, visual inspection of these aggregate interaction maps reveals that inter-sister interactions at these cohesin sites occur at shorter distances than intra-sister interactions. This becomes more evident when we calculate the difference between the two aggregate interaction maps (figure 4c). We observe in this difference plot an enrichment for inter-sister interactions over intra-sister interaction for cohesin sites separated by less than 20kb. Figure 4d illustrates this difference in another way by plotting the enrichment of inter and intra-sister signal over expected as a function of genomic distance from cohesin binding sites (figure 4d). Inter-sister interactions at cohesin sites preferentially occur at a distance up to 20kb, whereas, intra-sister interactions become more abundant at distances larger than 20kb. This can also be seen at an individual cohesin site (supplemental figure 10).

To explore and quantify these differences between inter-sister and intra-sister interactions in more detail, we analyzed pairwise cohesin-cohesin site interactions at different genomic distances. Interactions of cohesin sites within 10kb from each other are preferentially inter-sister interactions (figure 4a-c, second panel from left). This difference is also observed when cohesin sites are separated by 10-20kb (figure 4a-c, middle panel). Interestingly, this preference switches for pairwise cohesin interactions larger than 20kb: for cohesin sites separated by 20 to 35kb we observe a slight preference for intra-sister interactions (figure 4a-c, second panel from right). This difference for intra-sister interactions becomes much more prominent for pairwise cohesin interactions separated by 35 to 50kb (figure 4a-c, far right panel). Thus, cohesive cohesin enables interactions between sites at sister chromatids that are separated by only 5 to 25kb, whereas extruding cohesin generates loops along sister chromatids that can be as large as 50kb.

Above we described that findings in the SisterC data suggest that a given cohesin binding site can be engaged in an inter-sister interaction in one cell and an intra-sister interaction in another cell. To investigate this further, we leveraged the fact that these two types of interactions occur at different length scales. Specifically, we ranked SisterC signal of pairs of cohesin binding sites at different distances on their intensity (supplemental figure 11). We identified 284 cohesin binding sites that are engaged in the strongest inter-sister interactions (top 10 percent) with other cohesin binding sites that are located 10 to 20kb away. When we explore intra-sister interactions for this set of cohesin binding sites, we observed that these cohesin binding site pairs also display enriched intra-sister interactions 10 to 20kb away, albeit at lower frequency compared to their inter-sister cohesin-cohesin binding site interactions (supplemental figure 11a-c). Similarly, we identified 284 cohesin binding sites that mediate the strongest intra-sister interactions between cohesin sites separated by 35 to 50kb. These cohesin-cohesin binding site pairs also show enriched in inter-sister interactions, although at a lower frequency compared to intra-sister interaction signal (supplemental figure 11d-f). This suggests that there are cohesin sites that mediate both strong intra-sister interaction and strong inter-sister interactions. Further, we find an overlap of 65 cohesin binding sites) between cohesin binding sites that mediate strong inter-sister interactions with distal cohesin binding sites located at 10 to 20kb distance as well as strong intra-sister interactions with cohesin binding sites located at 35 to 50kb distance (supplemental figure 11g). This provides further support that a given cohesin binding site can engage in either inter-sister interactions or intra-sister interactions in different cells in the population.

## Discussion

Here we describe SisterC, a chromosome conformation capture technique that allows for detection of interactions between and along sister chromatids separately. SisterC leverages BrdU incorporation and single strand breaks induced by UV/Hoechst treatment, to assign Hi-C interactions as inter-sister or intra-sister interactions based on read orientation after sequencing.

First, SisterC reveals the extent to which sister chromatids are aligned. At centromeres the alignment is rather precise, possibly as a result of high density of cohesin binding sites that are engaged in inter-sister interactions at centromeric regions in every cell. The alignment is more loose along chromosome arms with inter-sister interactions spaced every 35kb on average. Along chromosome arms cohesin binding is observed every 10 to 15kb, however this result suggests that not every cohesin site will be bound in every cell or not every cohesin site will be engaged in inter-sister interactions. The alignment of sisters observed here resembles the pairing of homologues observed in *Drosophila*^25^. In that study, a similar analysis and comparison of *P*_inter_(*s*) and *P*_intra_(*s*) was used to infer the extent of alignment of homologues and regional variation in the precision of alignment chromosomes was observed along the length of the chromosomes. Although we see a clear difference in alignment between centromeric regions and chromosome arms in mitotic yeast sister chromatids, we do not observe different degrees of alignment along chromosome arms.

Second, we observe that inter-sister interactions likely mediated by cohesive cohesin, occur mostly between cohesin binding sites separated by less than 25kb. This distance is smaller than the distance between inter-sister interaction sites, which we estimated occur every 35kb on average (see above). One explanation for this can be that inter-sister interactions get established during S-phase relatively close to the replication fork that generates the sister chromatid pair^26–28^. Possibly, inter-sister interactions are initially very precise, but can possibly move to the closest cohesin binding site at convergent gene pairs on both sister chromatids^22,29,30^, producing a spacing that can be up to 25kb. Along these lines, we explored whether that this offset of inter-sister chromatid interactions would differ from genome-wide average at and around origins of replications. However, we find no clear differences in sister chromatid interactions along or between sister chromatids near origins of replication (supplemental figure 12). Interestingly, we do see a clear boundary at origins of replication in our control G1 Hi-C libraries (supplemental figure 12d).

Third, we observe that intra-sister interactions, possibly mediated by extruding cohesin, form larger loops ranging from 25 to 50kb. These cohesin-mediated extruded loops are established during G2/M-phase^7^. Interestingly, previous polymer simulation showed that yeast Hi-C data from mitotic cells is consistent with the formation of dynamic loops of around 35kb in size, which cover about 35% of the genome^7^. This is in agreement with our SisterC observations.

Cohesive cohesin and extruding cohesin are preferentially interacting at different genomic distances, mediate interactions independent from each other and are loaded and established in different phases in the cell cycle. This leads us to propose that these cohesin complexes are distinct, possibly having different subunit compositions or modifications. For instance, cohesin can be bound by either Scc2/4 or Pds5^31–33^, leading to cohesin complexes with different properties. Further, the acetylation status of a cohesin complex mediated by Eco1 is particularly important for establishment of cohesion^34,35^. Where in yeast both inter-sister interactions and intra-sister interactions in mitosis are mediated by cohesin complexes, in vertebrates these are formed by two different protein complexes: cohesin establishes sister chromatid cohesion and condensin I and II mediate intra-sister looping formation to compact chromosomes^7,36^. Additionally, it is important to note that in yeast a given cohesin binding site can be involved in both an inter-sister interaction and an intra-sister interaction, although most likely not occurring in the same cell at a given time. Due to a low cohesin binding density in yeast and absence of sequence specificity of cohesin complexes, there will be large cell-to-cell variation in which cohesin sites will be bound by either cohesive or extruding cohesins.

In the current SisterC procedure, selective depletion of DNA strands containing BrdU is not complete. Therefore, there is some level of cross contamination of inter- and intra-sister interactions. This level of contamination can be estimated by analyzing SisterC libraries of cells in G1-phase after one round of S-phase in the presence of BrdU. The reason why selective depletion is not complete is most likely related by the relatively low efficiency of DNA breakage after UV/Hoechst treatment. Importantly we did not identify particular types of molecules that are resistant to selective depletion. For instance, A-T content of the genomic site (supplemental figure 13a-b), distance from a digestion site (supplemental figure 13c-d) or regions near replication origins (supplemental figure 13e-f) do not affect assignment of interactions as being inter-sister or intra-sister.

SisterC allows the study of the significant topological challenge each cell faces during the cell cycle: the concordant chromosome compaction and sister chromatid separation during mitosis, particularly prophase. This process has been difficult to study by conventional Hi-C, because of its inability to distinguish between inter- and intra-sister interactions. We believe SisterC will have a broad applicability in different model organisms and during different phases of the cell cycle from late S-phase to mitosis.

## Methods

### Yeast synchronization and culture conditions

The YLV11 strain^18^ was used for all experiments in this study. For normal growth, cells were cultured in YP media with 2% galactose and 100μM thymidine at 30°C. Prior to synchronization, cultures were diluted to OD ~0.15 and allowed to double. To synchronize cells in late G1, 5 μM alpha factor mating pheromone (zymoresearch #Y1001) was added for 2.5-3 hours until cells started schmoo formation. SisterC DpnII R2 and R3 received a boost of 500uM thymidine or BrdU after 2 hours of alpha factor synchronization. G1 arrested cells were washed 3 times and released in prewarmed media containing 1mM BrdU or Thymidine. For mitotic arrested cells, 1% DMSO was added after release and 10μg/mL was added 30 minutes later. Mitotic cells were harvested approximately 4.5 hours after alpha factor release. For G1 arrested cells, the culture was released for 2 hours, followed by a second alpha factor arrest for 3 hours. Cells were washed and pelleted for genomic DNA extraction for HPLC detection and stored at −80°C until further processing. For Hi-C and SisterC, cells were fixed with 3% formaldehyde for 20 minutes at 30°C while in shaker incubator. Fixing was quenched by adding 2.5M glycine for an additional 5 minutes at 30°C. Cells were washed twice in MilliQ and pelleted cells were stored at −80°C till further processing. Throughout synchronization protocol, cells were washed and fixed in 95% ethanol for flow cytometry analysis.

### Flow cytometry

Ethanol fixed cells were resuspended and washed with 50mM NaCitrate. After mild sonication, cell walls were degraded with 10 units of zymolyase in PBS for 30 min at 30°C, followed by a lysis using 2M HCl and 0.5% Triton Tx-100 for 30 min at room temperature, followed by 30 min incubation at room temperature in 0.1M NaB_4_O_7_. Cells were then washed and resuspended in PBS, 1% milk, 0.2% Tween and 1:20 anti-BrdU-FITC (ThermoFisher #11-5071-42) and incubated for 30 min at RT. Cells were again washed and resuspended in PBS, 1% milk, 0.2% Tween, 0.25mg/mL RNase A and 10mg/mL propidium iodide and incubated at 37°C for 30 min. Cells were washed and resonicated before flow cytometry detection using a MACSquant flow cytometer. Data was analyzed using FlowJo software.

### Amplification of DNA fragments containing after UV/Hoechst treatment

DNA fragments (686 bp length) were amplified for 15 cycles using DreamTaq (ThermoFisher # EP1701) in presence of 100%, 90%, 50%, 10% or 0% BrdUTP, supplemented with dTTP. Amplified DNA fragments were incubated in TLE with 100ng/uL Hoechst 33342 (ThermoFisher #H3570) for 15 min at room temperature while protecting from light, followed by UV radiation at 2.7kJ/m^2^. Samples were washed 3 times in 30KDa amicon columns (MilliPore # UFC5030BK) and amplified with 10 PCR cycles.

### HPLC separation and LCMS analysis

Cells were harvested, pelleted and frozen at −80°C for HPLC analysis from 300mL yeast cultures at OD ~0.3 (approximately 600 million cells). Frozen cells were washed in 1mL spheroplasting buffer and lysed for 10 minutes at 35°C using 0.5% beta-mercaptoethanol and 10ug/mL zymolyase (Zymoresearch # E1005). Cells were washed in 1x NEBuffer 3.1 (NEB #B7203S) and incubated with proteinase K twice for 12 hours and an additional 2 hours at 65°C. DNA was extracted with phenol:chloroform, followed by ethanol precipitation and RNA digestion using RNase A. DNA samples were first washed with milliQ water using a 3 kDa cut-off Amicon filter. Digestion into nucleosides was carried out on 3 μg aliquots using ‘nucleoside digestion enzyme mix’ from New England Biolabs (NEB#M0649) and incubated overnight at 37°C. Once digested, aliquots from the same sample were pooled, filtered, and rinsed through a 3 kDa Amicon filter with milliQ water. The flow through was concentrated and quantified by A260 for subsequent HPLC injection. Digested deoxynucleosides were resolved using an Agilent 1260 Infinity HPLC with a Synergi C18 4-μm Fusion-RP 80Å 250 x 4.6 mm LC column. Nucleosides were resolved over 35 minutes using an isocratic gradient of 2-22% [95% acetonitrile, 5% 20 mM ammonium acetate pH 4.5] in [20 mM ammonium acetate] at 25°C, using a flow rate of 0.5 mL/min. This was followed by wash steps at higher eluent strengths between runs. Absorbance was recorded at 260 nm and 279 nm. Normal deoxynucleosides were quantitated using HPLC peak areas over three repeats, and published values for molar extinction coefficients at 260 nm [ɛ260 (M-1 cm-1) adenosine = 15400, cytidine = 7300, guanosine = 11700, thymidine = 8800^37^. The molar extinction coefficients for bromodeoxyuridine at 260 and 279 nm were empirically measured and found to be 9229 and 5003, respectively. For mass spectrometry analysis, deoxynucleosides were first resolved using the HPLC method above, and UV peaks were manually collected in separate vials. These were evaporated to dryness, resuspended in 50 μL milliQ water, and desalted on the same HPLC column using an isocratic gradient of 0-40% [98% acetonitrile, 2% water] in [90% water, 10% acetonitrile] over 15 min at 25°C, using a flow rate of 1 mL/min. The desalted peak was collected, evaporated to dryness, then resuspended in 20 μL milliQ water and analysed by high resolution liquid chromatography mass spectrometry. Mass spectrometry was carried out in positive ESI mode on an Agilent 6530 Accurate-Mass Q-TOF LC/MS linked to a pre-injection Agilent 1260 infinity HPLC. Samples were run in milliQ water containing 0.1 % formic acid. Mass spectrometry settings to detect nucleosides were as follows; gas temperature 350°C, nebulizer gas rate 45 psig, drying gas 10 L/min, VCap 4000 V, fragmentor voltage 120 V, skimmer voltage 65 V. 5 mM stock solutions of RNA nucleosides A, U, C, and G (ChemGenes) were made up in milliQ water. HPLC injections of digested deoxynucleosides were doped with the addition of 2 μL of each of the RNA nucleosides. Nucleosides were resolved using the standard separation method above.

### SisterC and Hi-C library preparation

For each mitotic SisterC or Hi-C library approximately 300 million cells or 100mL culture at OD ~0.3 was used. For each G1 SisterC or Hi-C library this was double, approximately 600 million cells or 100mL culture at OD ~0.6. There were 3 biological replicates produced using DpnII as restriction enzyme and one replicate using HindIII. SisterC and Hi-C were preformed according to previously published Hi-C protocol for yeast^23^, with several major modifications. Samples were split for Hi-C and SisterC library production before treatment with Hoechst/UV. Briefly, cells were fixed and stored as described above. Crosslinked cells were thawed, washed and resuspended in spheroplasting buffer (1M Sorbitol, 50mM Tris pH 7.5). Cells were lysed by addition of 0.5% beta-mercaptoethanol and 10ug/mL zymolyase (Zymoresearch # E1005) and incubated for 10 min at 35°C. Cells were washed twice with 1x NEBuffer 3.1 (for DpnII libraries) or NEBuffer 2.1 (for HindIII libraries). Chromatin was solubilized with 0.1% SDS for 10 minutes at 65°C, followed by quenching with 1% Triton X-100. Chromatin was digested with 400U HindIII or DpnII overnight at 37°C. After inactivation of restriction enzyme for 20 min at 65°C, DNA ends were filled in with nucleotides and supplemented with biotin-14-dCTP (LifeTech #19518018) for HindIII libraries and biotin-14-dATP (LifeTech #19524016) for DpnII libraries for 4 hours at 23°C. DNA fragments were ligated with T4 DNA ligase (LifeTech #15224090) for 4 hours at 16°C in reactions of 75 μL each. All ligation reactions were combined and samples were treated with proteinase K overnight at 65°C. DNA was purified using 1:1 phenol:chloroform and ethanol precipitation. Samples treated with RNAse A and biotin from unligated ends were removed using T4 DNA polymerase. DNA was sonicated and size-selected using AMpure XP beads (Bedman coulter #A63881) to obtain fragments sized 600-800bp. We performed end repair and a-tailing prior to illumina TruSeq adapter ligation. Each sample was split in two to obtain one SisterC library treated with UV and Hoechst and one Hi-C library without treatment from the same biological sample. SisterC libraries were treated in two reaction volumes of 50uL each with 100ng/uL Hoechst 33342 (ThermoFisher #H3570) for 15 min at room temperature while protecting from light, followed by UV radiation at 2.7kJ/m^2^. Samples were washed with TLE 3 times in 30KDa amicon columns (MilliPore # UFC5030BK). Both SisterC and Hi-C libraries were then enriched for biotin-containing fragments by pull down with MyOne Streptavidin C1 beads (Life Tech #65-001). Libraries were amplified, cleaned from pcr primers and sequenced using paired end 50bp reads on an Illumina HiSeq4000 platform. All libraries within a set of replicates were amplified with the same number of PCR cycles.

### SisterC and Hi-C analysis

Hi-C and SisterC FASTQ sequencing files were mapped to saccer3 yeast reference genome using publicly available distiller-nf mapping pipeline (https://github.com/mirnylab/distiller-nf) and downstream analysis tools pairtools (https://github.com/mirnylab/pairtools) and cooltools (https://github.com/mirnylab/cooltools). Briefly, reads were mapped with bwa-mem, deduplicated and filtered for mapping quality, resulting in only “valid reads”. Reads were classified as inter-sister reads when one read end was mapped as + orientation and the other end as − read orientation. Reads were classified as intra-sister reads when both read ends mapped as + or − read orientation. For downstream analysis, interactions at a shorter distance than 1500bp were removed. Interactions were binned at 1kb, 2kb and 10kb resolution using cooler^38^. Iterative balancing was applied to all matrices, individually for inter-sister and intra-sister interactions, while ignoring two bins from the diagonal^39^. Hi-C and SisterC statistics for all samples are provided in supplemental table 1.

Distance decays were plotted from valid pairs separated by read orientation were used to calculate contact frequency (P) as a function of genomic distance (s) using cooltools code. Pile up plots on genomic loci were produced using valid interactions binned at 1kb resolution separated for inter- or intra-sister interactions and contained only interactions at distances larger than 1500bp. Observed over expected values were calculated using expected files of matching conditions (e.g. expected file of inter-sister interactions of SisterC library for matching observed interactions of inter-sister interactions of SisterC library). Heatmaps were plotted using modified cooltools code. For centromere pile up plot, the directionality of the centromere DNA elements was taken into account. Anchor plots were plotted from valid interactions binned at 2kb resolution, which were separated for inter or intra-sister interactions and contained only interactions at distances larger than 1500bp.

### Publicly available datasets used in this manuscript

Scc1 calibrated ChIP-seq tracks from Hu et al^20^ were used for cohesin pile up SisterC heatmaps and ChIP-seq tracks in figure 1. This dataset is available on GEO under accession number GSM1712309. Peaks were called on this dataset using MACS2. Pairwise cohesin interactions were compiled by listing all possible pairwise combinations of cohesin peak sites in cis, followed by separation on distance between cohesin pairs (smaller than 10kb, 10 to 20kb, 20 to 35kb and 35 to 50kb). Cohesin sites in a 50kb window around centromeres and on all of chrXII and chrIV were removed from the dataset. Additionally Hi-C samples from cdc45 mutant cells were used from Schalbetter et al ^7^ to investigate distance decay. This dataset is available on GEO under accession number GSM2327664. This data was processed identical to Hi-C libraries produced for this study. Sites of origin of replication were downloaded from OriDB (http://www.oridb.org/)^40^.

### Code availability

Hi-C mapping pipeline distiller-nf is available on https://github.com/mirnylab/distiller-nf. Downstream analysis tools pairtools and cooltools are available through https://github.com/mirnylab/pairtools and https://github.com/mirnylab/cooltools.

## Acknowledgements

We thank all current and former Dekker lab members for helpful discussions, in particular Sergey V. Venev for advice on data analysis. We thank Jennifer Benanti, Heather Arsenault and Nick Rhind for advice on culturing and synchronizing yeast and providing the yeast strain used in this study. We thank Stephanie Schalbetter, Nicola Minchell and Matthew Naele for providing their yeast Hi-C protocol and suggestions. We also thank Bas van Steensel for advice and suggestions. This work was supported by grants from the National Institutes of Health (HG003143 to J.D. and NS111990 to J.K.W.). J.D. is an investigator of the Howard Hughes Medical Institute.

## Author contributions

M.E.O. and J.D. conceived and designed the project. M.E.O. cultured yeast, generated and analyzed SisterC and Hi-C datasets and performed flow cytometry experiments. A.K.H. and J.K.W. designed HPLC and mass spectrometry experiments, A.K.H. performed and analyzed HPLC and MS experiments. M.E.O and J.D. wrote the manuscript.

## Competing interest statement

All authors declare no conflict of interest.

## Data access

All genomic data generated for this study will be available on the NCBI Gene Expression Omnibus upon publication. Code used for mapping and analysis of Hi-C data is available on github (https://github.com/mirnylab/distiller-nf and https://github.com/mirnylab/cooltools).

