## Supplemental Materials for "Detecting chromatin interactions along and between sister chromatids with SisterC"

### Supplemental Materials – Table of Contents

#### Supplemental Figures

Supplemental figure 1 – Outline of SisterC culture conditions

Supplemental figure 2 – PCR amplification of BrdU containing DNA fragments with and without Hoechst and UV treatment

Supplemental figure 3 – FACS data of all SisterC synchronization experiments

Supplemental figure 4 – HPLC nucleoside digestion data

Supplemental figure 5 – Definitive assignment of nucleoside peaks

Supplemental figure 6 – Distance decay plots of all SisterC replicates

Supplemental figure 7 – Distance decay plots of all SisterC control experiments

Supplemental figure 8 – Observed/expected interaction heatmap of chrXIII:450,000-550,000

Supplemental figure 9 – Centromeric region of chrXII

Supplemental figure 10 – Anchor plot on ChrXIII:469000-469001

Supplemental figure 11 – Ranking of SisterC signal on pairs of cohesin binding sites at different genomic distances

Supplemental figure 12 – SisterC pile up plots of origins of replication

Supplemental figure 13 – Efficiency of SisterC inter- and intra-sister interaction assignment

#### Supplemental Table

Supplemental Table 1 – SisterC and Hi-C mapping statistics

a

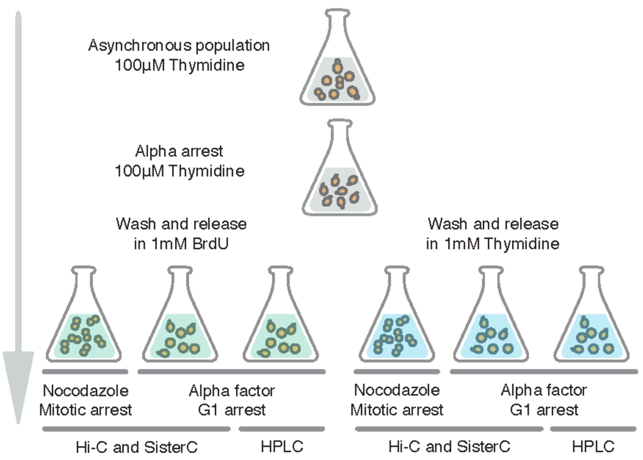


**Supplemental figure 1 – Outline of SisterC yeast culture conditions. (a)** Asynchronous yeast cultures are cultured and synchronized in late G1 using alpha factor. Cells are released in media containing BrdU or Thymidine, followed by an arrest in mitosis using nocodazole or G1 using a second alpha factor arrest. Cells are harvested and prepared for Hi-C or SisterC library production or processed for BrdU detection using HPLC.

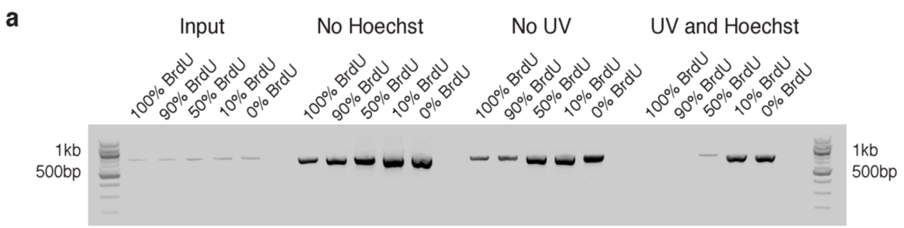


**Supplemental figure 2 – Depletion of BrdU containing DNA molecules by PCR.** DNA fragments were amplified in presence of 0%,10%,50%,90% and 100% BrdU to allow for incorporation in both strands (first 5 lanes). This was followed by treatment of UV only (second 5 lanes), Hoechst only (third 5 lanes) or treatment with both UV and Hoechst (last 5 lanes). Fragments containing more than 10% BrdU did not get amplified after UV/Hoechst treatment.

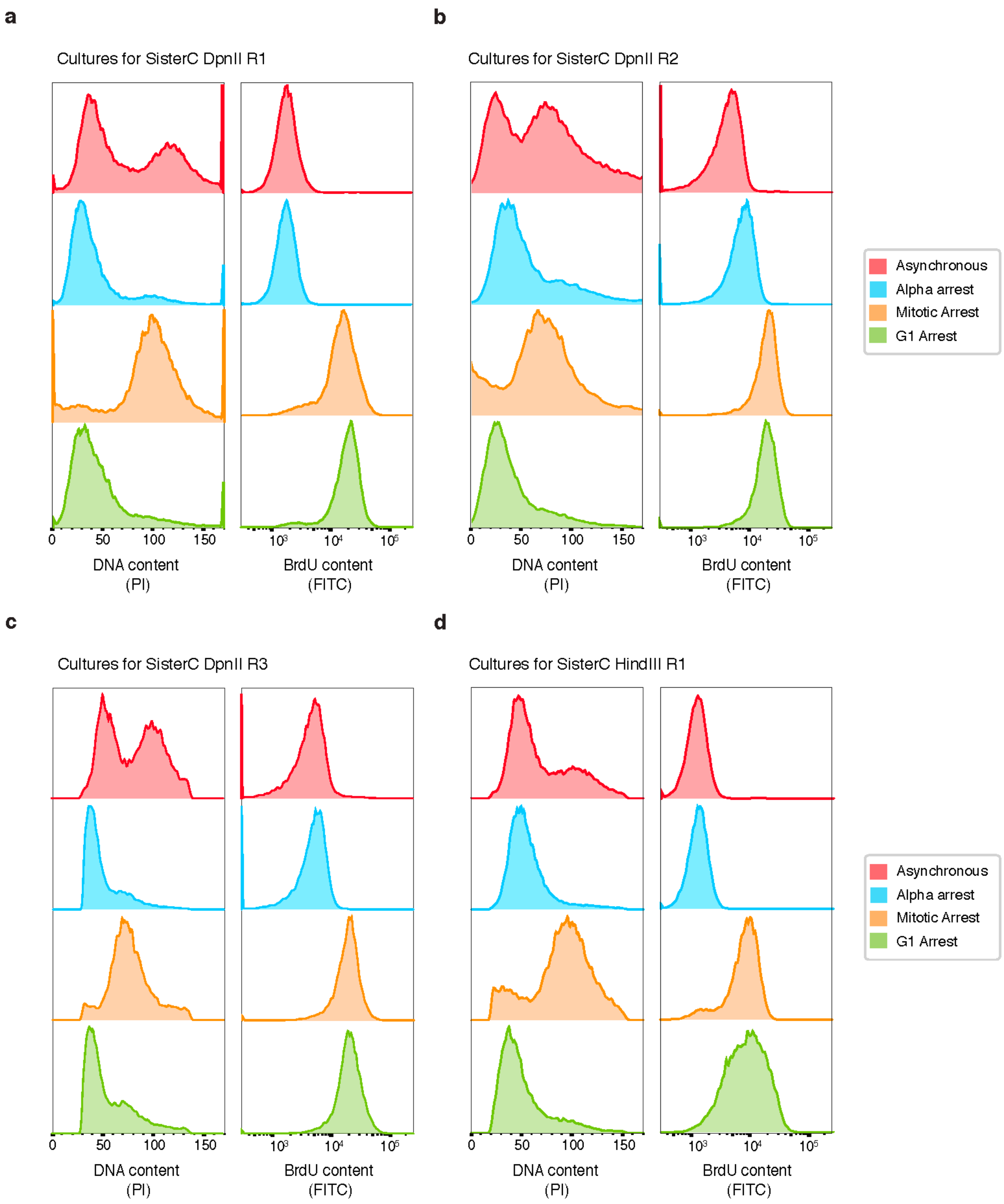


**Supplemental figure 3 – FACS data of all SisterC experiments.** Flow cytometry data detecting cell cycle profile and BrdU content of cultures grown for SisterC DpnII replicate 1 (**a**), SisterC DpnII replicate 2 (**b**), SisterC DpnII replicate 3 (**c**) and HindIII R1 (**d**).

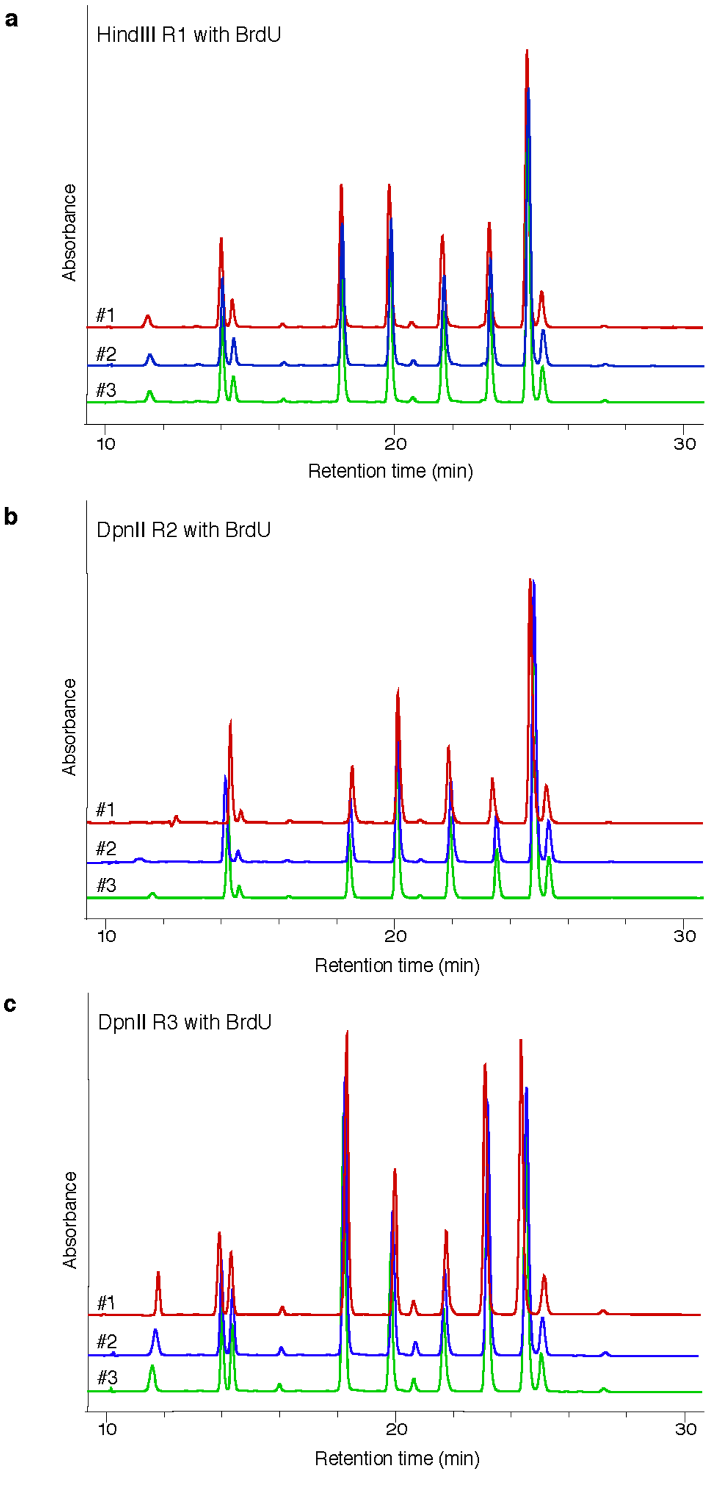


**Supplemental figure 4 – HPLC Nucleoside Digestion data.** Stacked HPLC traces from 3 repeats of each SisterC library HindIII R1 (**a**), DpnII R2 (**b**) and DpnII R3 (**c**).

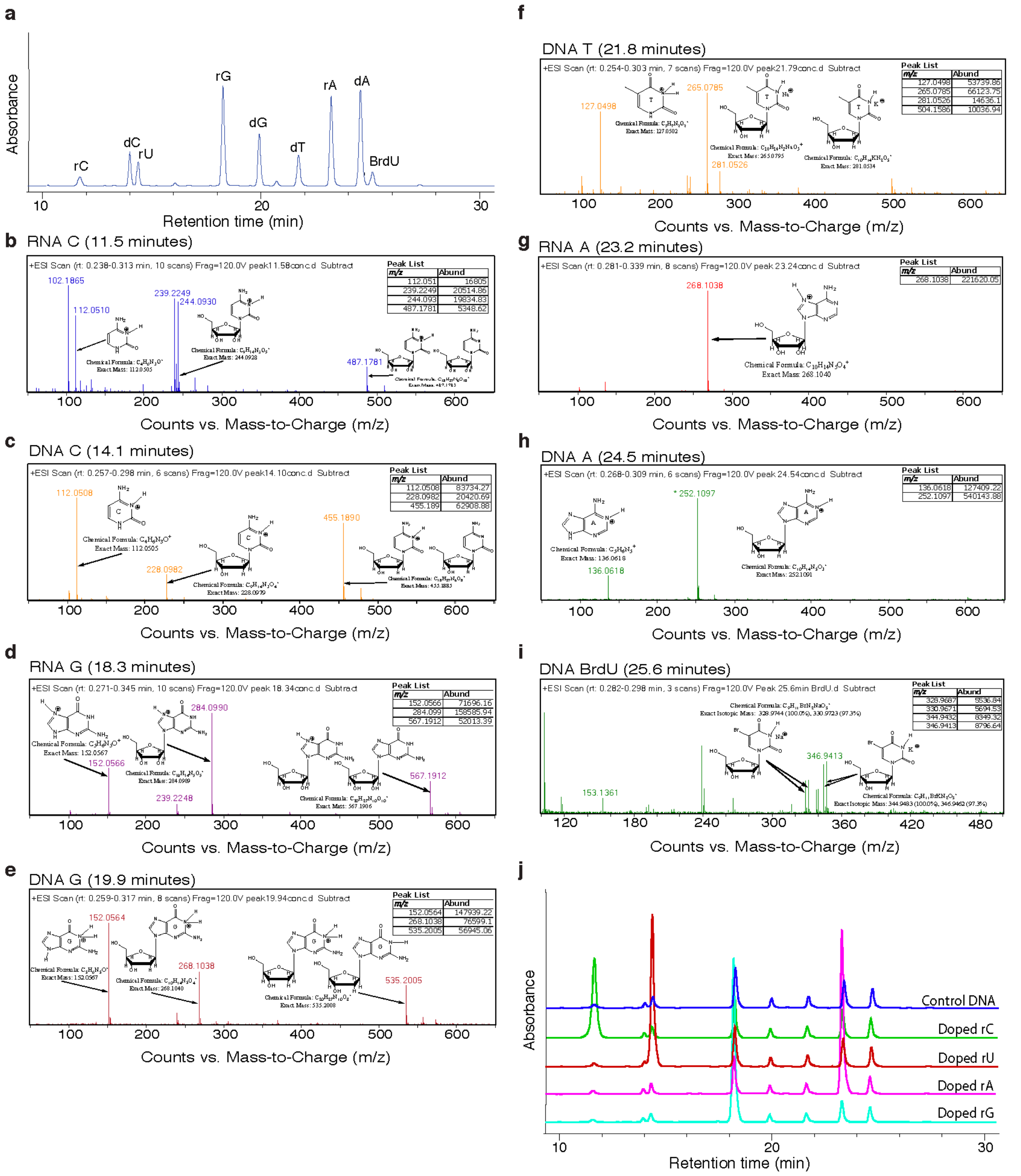


**Supplemental figure 5 – Definitive assignment of nucleoside peaks.** (**a**) HPLC trace showing peaks from a digested, BrdU containing sample of DNA; provided for reference of the peaks assigned in the following sections. (**b-i**) Mass spectrometry analysis of individual peaks shown in panel a. No diagnostic m/z peaks were found for the 14.4 minute peak, however RNA U doping (brown trace, panel **j**) was used to unambiguously assign this peak as RNA U. RNA A has the same chemical formula and mass as dG (218.1040 g/mol). Both the fragmentation pattern of the DNA G peak (panel e) and the RNA A doping experiment (magenta trace panel j) were used to confirm the assignment of the 23.2 minute peak as RNA A (panel g). (**j**) Stacked HPLC runs of digested, non-BrdU-treated control DNA doped with each ribonucleoside individually (2 uL of 5 mM solution of rU, rC, rA or rG was added to the digested DNA immediately before HPLC analysis).

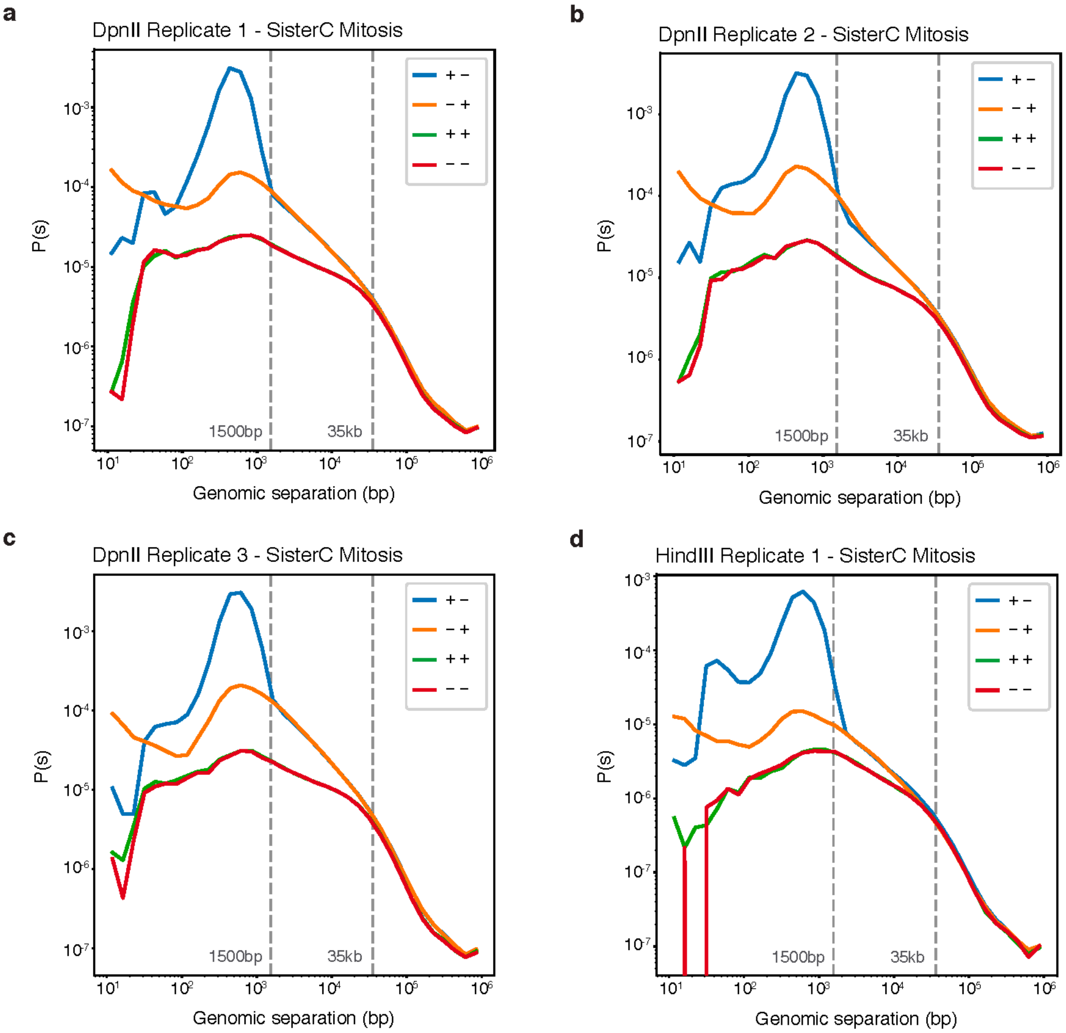


**Supplemental figure 6 – Distance decay plots of all SisterC replicates.** Distance decay plots of all SisterC mitotic libraries: DpnII replicate 1 (**a**), DpnII replicate 2 (**b**), DpnII replicate 3 (**c**) and HindIII replicate 1 (**d**)

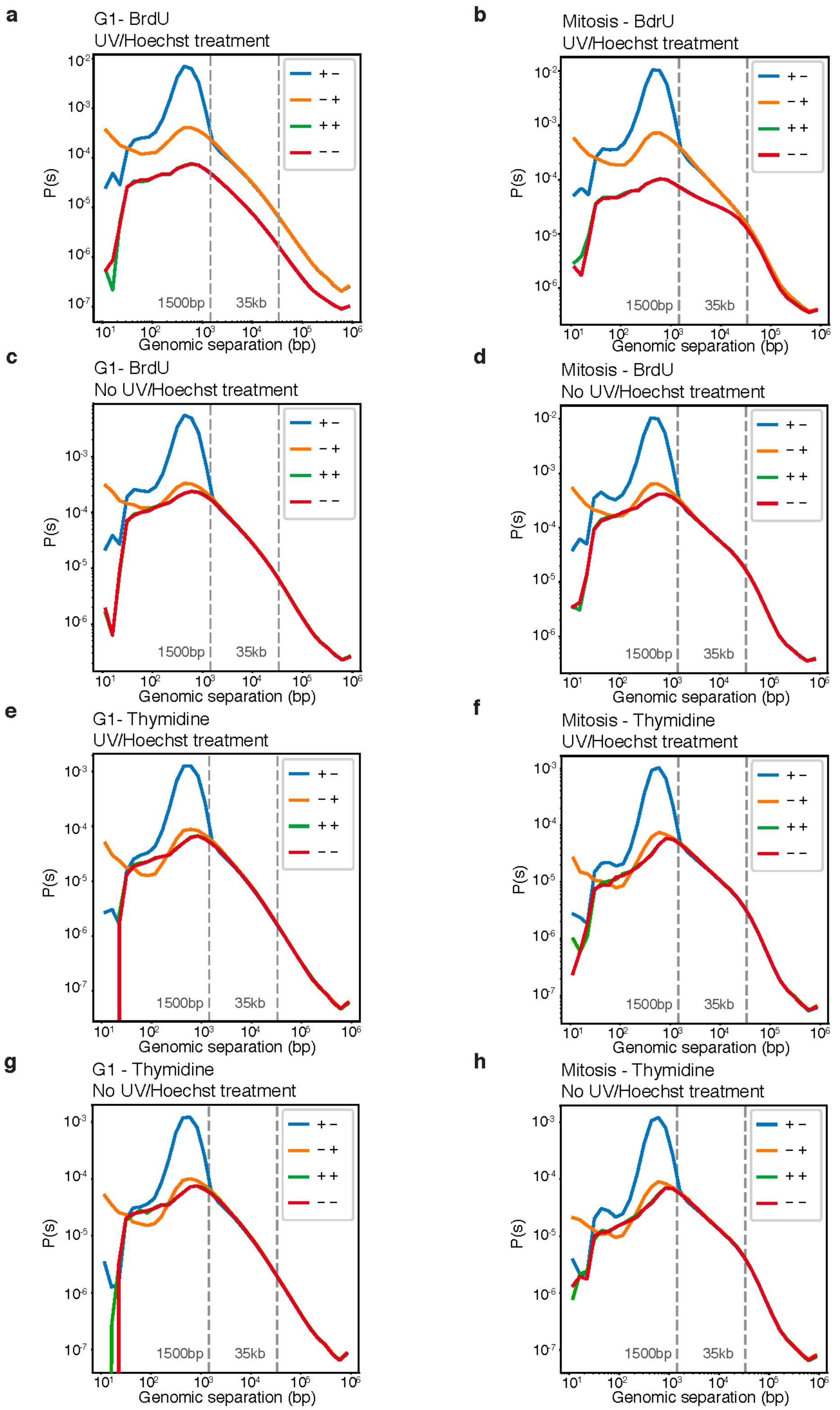


**Supplemental figure 7 – Distance decay plots of all SisterC control experiments.** Distance decay plots of all SisterC control libraries: (**a-b**) G1 (a) and mitotic (b) arrested cells grown in BrdU and treated with UV/Hoechst. (**c-d**) G1 (c) and mitotic (d) arrested cells grown in BrdU, not treated with UV/Hoechst. (**e-f**) G1 (e) and mitotic (f) arrested cells grown in Thymidine, treated with UV/Hoechst. (**g-h**) G1 (g) and mitotic (h) arrested cells grown in Thymidine, not treated with UV/Hoechst.

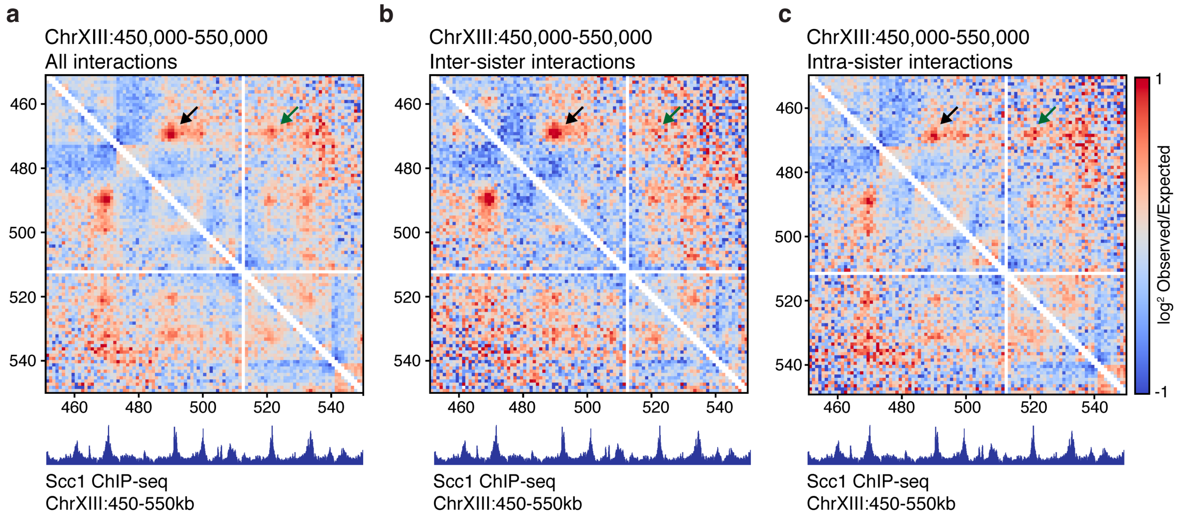


**Supplemental figure 8 – Observed/expected interaction heatmap of chrXIII:450,000-550,000.** Log2 observed over expected interaction frequency on region ChrXIII:450,000-550,000 (as shown in main figure 2) for all interactions (**a**), inter-sister interactions (**b**) and intra-sister interactions (**c**).

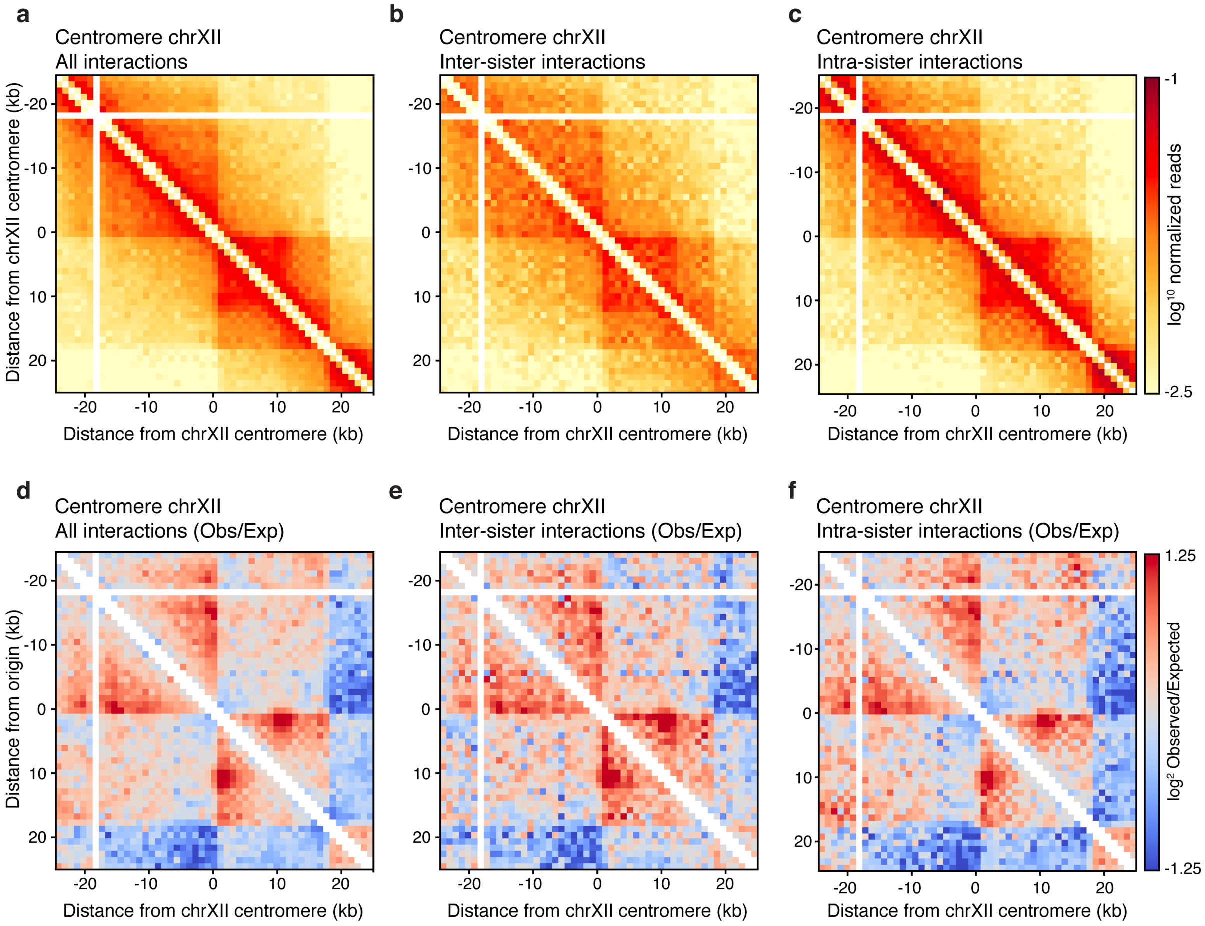


**Supplemental figure 9 – Centromeric region of chrXII. (a-c)** Interaction heatmaps of 50kb window around the centromeric region of chrXII, for all interactions (a), inter-sister interactions (b) and intra-sister interactions (c). (**d-f**) Log2 observed over expected interaction heatmaps of all interactions (d), inter-sister interactions (e) and intra-sister interactions (f) on the centromeric region on chrXII.

**a**

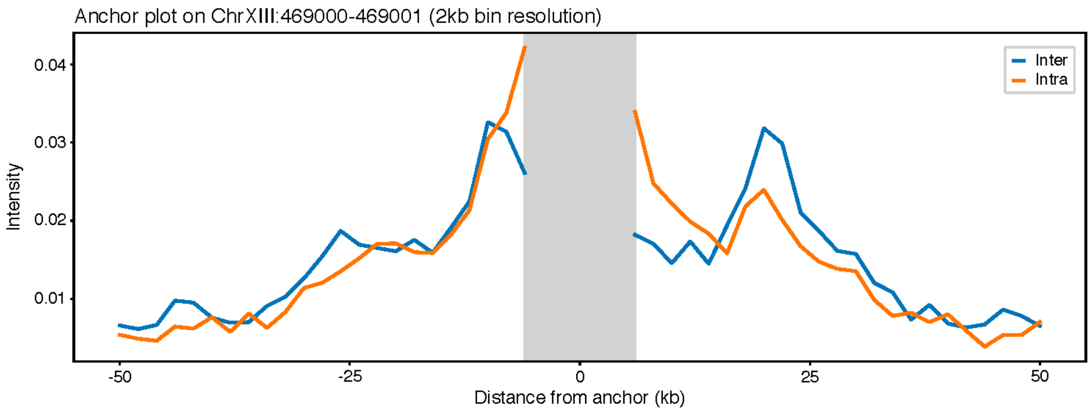


**Supplemental figure 10 – Anchor plot on ChrXIII:469000.** Interaction frequency of inter-sister and intra-sister interactions anchored on chrXIII:469,000 shows higher frequency of inter-sister interactions at 25kb distance from the anchor.

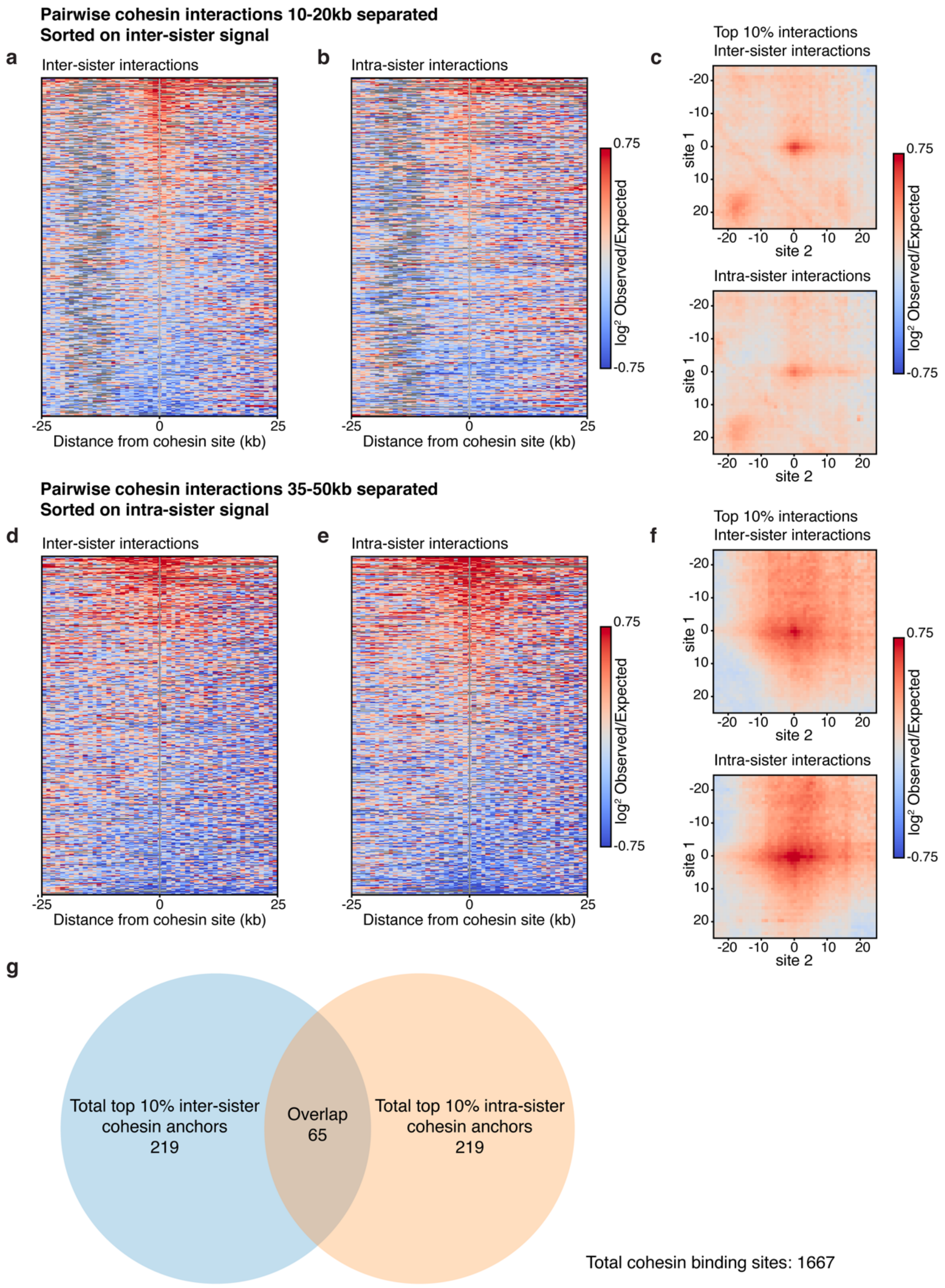


**Supplemental figure 11 – Ranking of SisterC signal on pairs of cohesin binding sites at different genomic distances. (a-b)** Inter-sister interaction (a) and intra-sister interaction (b) frequency of pairs of cohesin binding sites separated by 10 to 20kb were ranked by inter-sister interaction intensity for 10kb window around the cohesin binding site. (**c**) Interaction pile up plots of inter-sister and intra-sister interactions of the top 10 percent sites that were ranked in (a). (**d-e)** Inter-sister interaction (d) and intra-sister interaction (e) frequency of pairs of cohesin binding sites separated by 35 to 50kb were ranked by intra-sister interaction intensity for 10kb window around the cohesin binding site. (**f**) Interaction pile up plots of inter-sister and intra-sister interactions of the top 10 percent sites that were ranked in (e). (**g**) 284 cohesin sites that anchor the top 10% inter-sister interacting cohesin pairs at 10 to 20kb distance were identified as well as again 284 cohesin sites that anchor the top 10% intra-sister interacting cohesin pairs at 35 to 50kb distance. 65 cohesin sites are found to mediate both top 10% inter-sister cohesin-cohesin interactions and top 10% intra-sister cohesin-cohesin interactions.

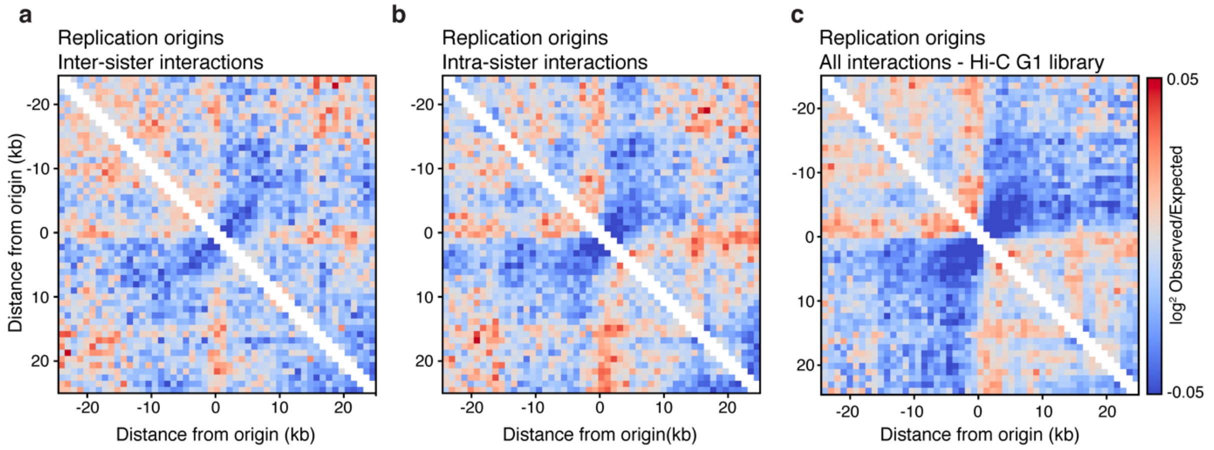


**Supplemental figure 12 – SisterC pile up plots of origins of replication. (a-b)** Mitotic SisterC inter-sister interactions (a) and intra-sister interactions (b) plotted on replication origins. (**c**) Pile up plot of Hi-C data of G1 alpha arrested cells on replication origins.

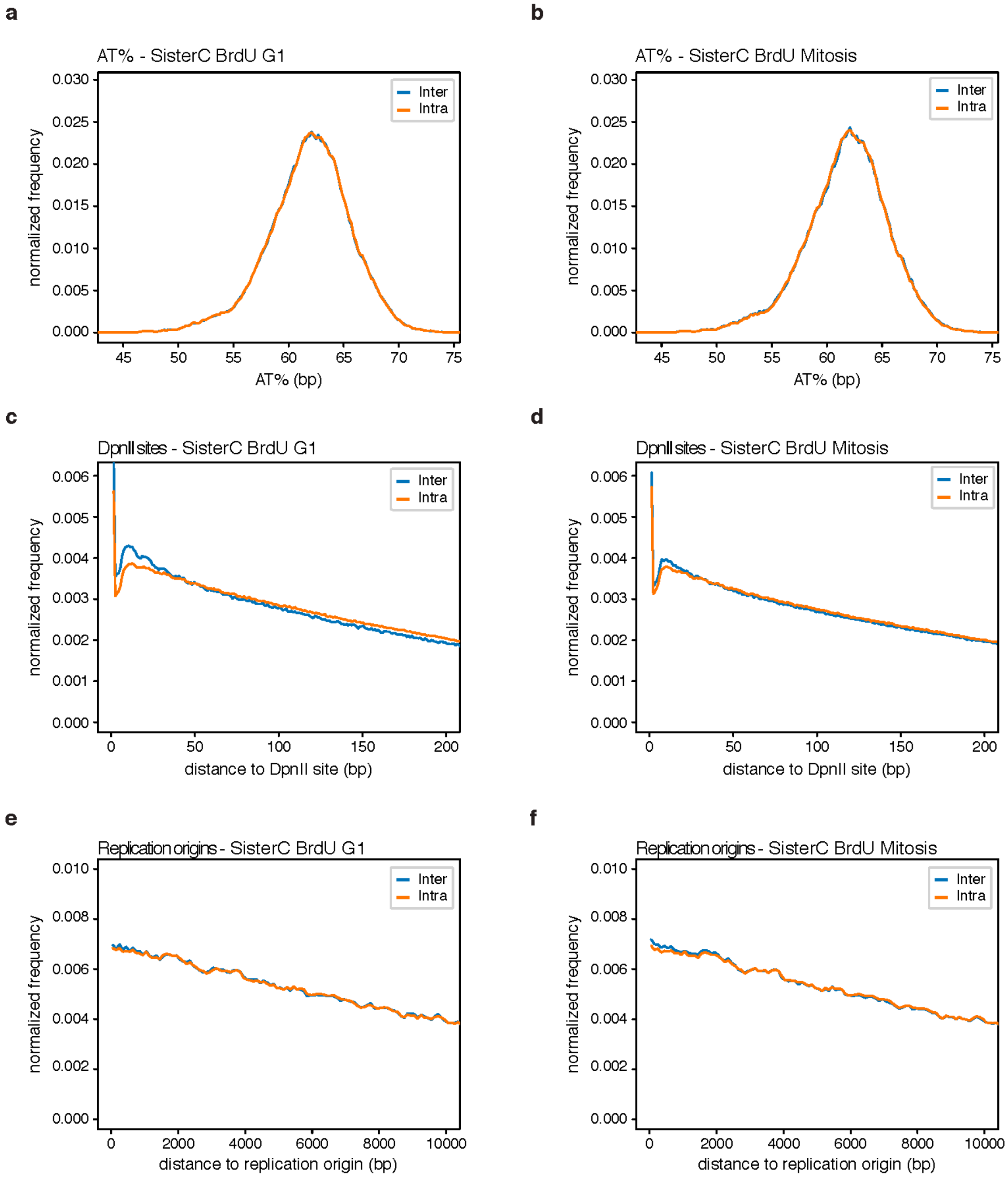


**Supplemental figure 13 – Efficiency of SisterC inter- and intra-sister interaction assignment.** (**a-b**) Normalized frequency SisterC G1 (a) and mitotic (b) inter-sister and intra-sister interactions as a function of AT percentage. (**c-d**) Normalized frequency SisterC G1 (c) and mitotic (d) inter-sister and intra-sister interactions as a function of distance to DpnII digestion site. (**e-f**) Normalized frequency SisterC G1 (e) and mitotic (f) inter-sister and intra-sister interactions as a function of distance to origins of replication.

| Replicate | Release in | Synchronized | UV/Hoechst treatment | Reads total | Total Mapped | | Total valid reads (no duplicates) | | Cis all interactions | | Cis all >1500bp | | Cis intra-sister >1500bp | | Cis inter-sister >1500bp | | Trans all | | Trans intra-sister | | Trans inter-sister | |
|---|---|---|---|---|---|---|---|---|---|---|---|---|---|---|---|---|---|---|---|---|---|---|
| | | | | # | # | % of total | # | % of mapped | # | % of valid | # | % of cis | # | % of cis >1500 | # | % of cis >1500 | # | % of valid | # | % of trans | # | % of trans |
| **DpnII R1** | BrdU | Mitosis | Yes | 2.27E+07 | 1.57E+07 | 69.11 | 1.42E+07 | 90.71 | 1.04E+07 | 73.32 | 5.12E+06 | 49.16 | 3.20E+06 | 62.42 | 1.92E+06 | 37.58 | 3.79E+06 | 26.68 | 1.90E+06 | 50.00 | 1.90E+06 | 50.00 |
| | BrdU | Mitosis | No | 2.28E+07 | 1.54E+07 | 67.33 | 1.57E+07 | 101.97 | 1.11E+07 | 70.64 | 6.78E+06 | 61.30 | 3.40E+06 | 50.11 | 3.38E+06 | 49.89 | 4.60E+06 | 29.36 | 2.30E+06 | 50.02 | 2.30E+06 | 49.98 |
| | BrdU | G1 | Yes | 2.27E+07 | 1.57E+07 | 69.11 | 1.46E+07 | 92.99 | 9.50E+06 | 65.20 | 3.96E+06 | 41.71 | 3.12E+06 | 78.90 | 8.36E+05 | 21.10 | 5.07E+06 | 34.80 | 2.53E+06 | 49.99 | 2.53E+06 | 50.01 |
| | BrdU | G1 | No | 2.28E+07 | 1.54E+07 | 67.33 | 1.46E+07 | 95.10 | 8.65E+06 | 59.19 | 4.60E+06 | 53.14 | 2.31E+06 | 50.24 | 2.29E+06 | 49.76 | 5.96E+06 | 40.81 | 2.98E+06 | 49.99 | 2.98E+06 | 50.01 |
| | Thymidine | Mitosis | Yes | 2.46E+07 | 1.66E+07 | 67.55 | 1.68E+07 | 101.09 | 1.27E+07 | 75.30 | 8.29E+06 | 65.55 | 4.16E+06 | 50.15 | 4.13E+06 | 49.85 | 4.15E+06 | 24.70 | 2.08E+06 | 50.03 | 2.07E+06 | 49.97 |
| | Thymidine | Mitosis | No | 2.43E+07 | 1.70E+07 | 69.93 | 1.60E+07 | 94.33 | 1.21E+07 | 75.21 | 8.29E+06 | 65.55 | 4.16E+06 | 50.15 | 4.13E+06 | 49.85 | 3.98E+06 | 24.79 | 1.99E+06 | 49.99 | 1.99E+06 | 50.01 |
| | Thymidine | G1 | Yes | 2.46E+07 | 1.66E+07 | 67.55 | 1.56E+07 | 93.68 | 8.78E+06 | 56.42 | 4.50E+06 | 51.20 | 2.26E+06 | 50.16 | 2.24E+06 | 49.84 | 6.78E+06 | 43.58 | 3.39E+06 | 50.00 | 3.39E+06 | 50.00 |
| | Thymidine | G1 | No | 2.04E+07 | 1.37E+07 | 67.15 | 1.30E+07 | 94.82 | 7.19E+06 | 55.36 | 3.82E+06 | 53.08 | 1.92E+06 | 50.26 | 1.90E+06 | 49.74 | 5.80E+06 | 44.64 | 2.90E+06 | 50.03 | 2.90E+06 | 49.97 |
| **DpnII R2** | BrdU | Mitosis | Yes | 1.63E+08 | 1.13E+08 | 69.37 | 7.79E+07 | 69.03 | 5.04E+07 | 64.67 | 2.22E+07 | 44.07 | 1.37E+07 | 61.86 | 8.47E+06 | 38.14 | 2.75E+07 | 35.33 | 1.38E+07 | 50.01 | 1.38E+07 | 49.99 |
| | BrdU | Mitosis | No | 1.64E+08 | 1.14E+08 | 69.32 | 8.88E+07 | 78.21 | 5.23E+07 | 58.84 | 2.95E+07 | 56.47 | 1.48E+07 | 50.31 | 1.47E+07 | 49.69 | 3.66E+07 | 41.16 | 1.83E+07 | 49.99 | 1.83E+07 | 50.01 |
| | BrdU | G1 | Yes | 1.16E+08 | 8.11E+07 | 69.87 | 5.16E+07 | 63.64 | 3.19E+07 | 61.90 | 1.03E+07 | 32.30 | 8.24E+06 | 79.90 | 2.07E+06 | 20.10 | 1.97E+07 | 38.10 | 9.83E+06 | 50.00 | 9.83E+06 | 50.00 |
| | BrdU | G1 | No | 1.23E+08 | 8.43E+07 | 68.49 | 6.35E+07 | 75.40 | 3.23E+07 | 50.83 | 1.48E+07 | 45.86 | 7.46E+06 | 50.36 | 7.35E+06 | 49.64 | 3.12E+07 | 49.17 | 1.56E+07 | 49.99 | 1.56E+07 | 50.01 |
| | Thymidine | Mitosis | Yes | 5.50E+07 | 3.72E+07 | 67.74 | 2.89E+07 | 77.52 | 1.67E+07 | 57.74 | 9.51E+06 | 57.04 | 4.80E+06 | 50.47 | 4.71E+06 | 49.53 | 1.22E+07 | 42.26 | 6.10E+06 | 50.00 | 6.10E+06 | 50.00 |
| | Thymidine | Mitosis | No | 5.45E+07 | 3.73E+07 | 68.46 | 2.91E+07 | 78.05 | 1.65E+07 | 56.58 | 9.97E+06 | 60.53 | 5.03E+06 | 50.49 | 4.93E+06 | 49.51 | 1.26E+07 | 43.42 | 6.32E+06 | 50.01 | 6.32E+06 | 49.99 |
| | Thymidine | G1 | Yes | 7.33E+07 | 5.03E+07 | 68.61 | 3.27E+07 | 65.12 | 1.57E+07 | 47.86 | 9.51E+06 | 57.04 | 4.80E+06 | 50.47 | 4.71E+06 | 49.53 | 1.71E+07 | 52.14 | 8.53E+06 | 50.01 | 8.53E+06 | 49.99 |
| | Thymidine | G1 | No | 5.90E+07 | 4.04E+07 | 68.47 | 3.22E+07 | 79.69 | 1.56E+07 | 48.42 | 9.97E+06 | 60.53 | 5.03E+06 | 50.49 | 4.93E+06 | 49.51 | 1.66E+07 | 51.58 | 8.30E+06 | 49.99 | 8.30E+06 | 50.01 |
| **DpnII R3** | BrdU | Mitosis | Yes | 1.88E+08 | 1.30E+08 | 69.13 | 7.57E+07 | 58.18 | 5.97E+07 | 78.84 | 3.06E+07 | 51.23 | 2.02E+07 | 65.94 | 1.04E+07 | 34.06 | 1.60E+07 | 21.16 | 8.01E+06 | 50.01 | 8.01E+06 | 49.99 |
| | BrdU | Mitosis | No | 1.86E+08 | 1.30E+08 | 69.80 | 1.07E+08 | 82.59 | 8.16E+07 | 76.12 | 5.15E+07 | 63.14 | 2.58E+07 | 50.15 | 2.57E+07 | 49.85 | 2.56E+07 | 23.88 | 1.28E+07 | 50.01 | 1.28E+07 | 50.01 |
| | BrdU | G1 | Yes | 1.54E+08 | 1.06E+08 | 68.46 | 8.52E+07 | 80.76 | 5.75E+07 | 67.52 | 2.40E+07 | 41.70 | 1.90E+07 | 79.02 | 5.03E+06 | 20.98 | 2.77E+07 | 32.48 | 1.38E+07 | 50.01 | 1.38E+07 | 49.99 |
| | BrdU | G1 | No | 1.89E+08 | 1.28E+08 | 67.85 | 1.10E+08 | 85.55 | 6.72E+07 | 61.25 | 3.58E+07 | 53.24 | 1.79E+07 | 50.19 | 1.78E+07 | 49.81 | 4.25E+07 | 38.75 | 2.12E+07 | 50.01 | 2.12E+07 | 49.99 |
| | Thymidine | Mitosis | Yes | 8.38E+07 | 5.57E+07 | 66.51 | 4.65E+07 | 83.55 | 3.48E+07 | 74.81 | 2.26E+07 | 64.82 | 1.13E+07 | 50.09 | 1.13E+07 | 49.91 | 1.17E+07 | 25.19 | 5.86E+06 | 50.01 | 5.86E+06 | 49.99 |
| | Thymidine | Mitosis | No | 8.92E+07 | 5.89E+07 | 66.05 | 5.05E+07 | 85.67 | 3.78E+07 | 74.86 | 2.47E+07 | 65.51 | 1.24E+07 | 50.05 | 1.24E+07 | 49.95 | 1.27E+07 | 25.14 | 6.34E+06 | 49.98 | 6.35E+06 | 50.02 |
| | Thymidine | G1 | Yes | 8.94E+07 | 5.87E+07 | 65.64 | 4.95E+07 | 84.32 | 2.95E+07 | 59.67 | 1.52E+07 | 51.45 | 7.64E+06 | 50.29 | 7.55E+06 | 49.71 | 2.00E+07 | 40.33 | 9.98E+06 | 50.01 | 9.97E+06 | 49.99 |
| | Thymidine | G1 | No | 9.65E+07 | 6.25E+07 | 64.77 | 5.26E+07 | 84.19 | 3.12E+07 | 59.34 | 1.68E+07 | 53.68 | 8.40E+06 | 50.14 | 8.36E+06 | 49.86 | 2.14E+07 | 40.66 | 1.07E+07 | 49.99 | 1.07E+07 | 50.01 |
| **HindIII R1** | BrdU | Mitosis | Yes | 5.81E+07 | 3.05E+07 | 52.48 | 1.08E+07 | 35.52 | 9.33E+06 | 86.20 | 3.45E+06 | 36.91 | 2.00E+06 | 58.13 | 1.44E+06 | 41.87 | 1.49E+06 | 13.80 | 7.47E+05 | 50.00 | 7.47E+05 | 50.00 |
| | BrdU | Mitosis | No | 3.90E+07 | 2.33E+07 | 59.79 | 1.63E+07 | 69.75 | 1.37E+07 | 84.20 | 6.46E+06 | 47.17 | 3.22E+06 | 49.84 | 3.24E+06 | 50.16 | 2.57E+06 | 15.80 | 1.29E+06 | 50.02 | 1.29E+06 | 49.98 |
| | BrdU | G1 | Yes | 8.04E+07 | 5.55E+07 | 69.04 | 2.74E+07 | 49.37 | 2.17E+07 | 79.09 | 5.63E+06 | 25.99 | 4.23E+06 | 75.15 | 1.40E+06 | 24.85 | 5.73E+06 | 20.91 | 2.86E+06 | 49.99 | 2.86E+06 | 50.01 |
| | BrdU | G1 | No | 4.29E+07 | 2.92E+07 | 68.20 | 2.40E+07 | 81.95 | 1.84E+07 | 76.93 | 5.68E+06 | 30.79 | 2.83E+06 | 49.81 | 2.85E+06 | 50.19 | 5.53E+06 | 23.07 | 2.76E+06 | 50.00 | 2.76E+06 | 50.00 |
| | Thymidine | Mitosis | Yes | 4.86E+07 | 2.66E+07 | 54.73 | 1.09E+07 | 41.10 | 9.31E+06 | 85.11 | 4.11E+06 | 44.20 | 2.09E+06 | 50.81 | 2.02E+06 | 49.19 | 1.63E+06 | 14.89 | 8.14E+05 | 50.00 | 8.14E+05 | 50.00 |
| | Thymidine | Mitosis | No | 5.34E+07 | 3.23E+07 | 60.59 | 1.31E+07 | 40.39 | 1.11E+07 | 85.03 | 4.89E+06 | 44.01 | 2.53E+06 | 51.71 | 2.36E+06 | 48.29 | 1.95E+06 | 14.97 | 9.78E+05 | 50.04 | 9.77E+05 | 49.96 |
| | Thymidine | G1 | Yes | 7.64E+07 | 4.59E+07 | 60.03 | 2.28E+07 | 49.66 | 1.68E+07 | 73.72 | 5.62E+06 | 33.47 | 2.89E+06 | 51.48 | 2.73E+06 | 48.52 | 5.99E+06 | 26.28 | 3.00E+06 | 50.02 | 2.99E+06 | 49.98 |
| | Thymidine | G1 | No | 8.84E+07 | 5.45E+07 | 61.66 | 3.39E+07 | 62.30 | 2.48E+07 | 73.00 | 8.60E+06 | 34.69 | 4.43E+06 | 51.55 | 4.16E+06 | 48.45 | 9.16E+06 | 27.00 | 4.59E+06 | 50.04 | 4.58E+06 | 49.96 |
